# Hydrogel-based slow release of a receptor-binding domain subunit vaccine elicits neutralizing antibody responses against SARS-CoV-2

**DOI:** 10.1101/2021.03.31.437792

**Authors:** Emily C. Gale, Abigail E. Powell, Gillie A. Roth, Emily L. Meany, Jerry Yan, Ben S. Ou, Abigail K. Grosskopf, Julia Adamska, Vittoria C. T. M. Picece, Andrea I. d’Aquino, Bali Pulendran, Peter S. Kim, Eric A. Appel

## Abstract

The development of effective vaccines that can be rapidly manufactured and distributed worldwide is necessary to mitigate the devastating health and economic impacts of pandemics like COVID-19. The receptor-binding domain (RBD) of the SARS-CoV-2 spike protein, which mediates host cell entry of the virus, is an appealing antigen for subunit vaccines because it is efficient to manufacture, highly stable, and a target for neutralizing antibodies. Unfortunately, RBD is poorly immunogenic. While most subunit vaccines are commonly formulated with adjuvants to enhance their immunogenicity, we found that clinically-relevant adjuvants Alum, AddaVax, and CpG/Alum were unable to elicit neutralizing responses following a prime-boost immunization. Here we show that sustained delivery of an RBD subunit vaccine comprising CpG/Alum adjuvant in an injectable polymer-nanoparticle (PNP) hydrogel elicited potent anti-RBD and anti-spike antibody titers, providing broader protection against SARS-CoV-2 variants of concern compared to bolus administration of the same vaccine and vaccines comprising other clinically-relevant adjuvant systems. Notably, a SARS-CoV-2 spike-pseudotyped lentivirus neutralization assay revealed that hydrogel-based vaccines elicited potent neutralizing responses when bolus vaccines did not. Together, these results suggest that slow delivery of RBD subunit vaccines with PNP hydrogels can significantly enhance the immunogenicity of RBD and induce neutralizing humoral immunity.

## 1. Introduction

The COVID-19 pandemic has had devastating health and economic impacts globally since SARS-CoV-2 first infected humans in late 2019. To date, COVID-19 has caused over 3.5 million deaths globally, including over 580,000 deaths in the US alone.^[1]^ Although behavioral and contact tracing interventions have slowed the spread and vaccines are becoming available in some regions, case numbers remain high in many parts of the world. Continued spread of SARS-CoV-2 will continue to be particularly harmful in regions that have limited resources and access to healthcare. High rates of asymptomatic transmission and the lack of effective treatments has made the virus difficult to contain.^[2]^ Deployment of effective vaccines is therefore a critical global health priority toward ending the COVID-19 pandemic. Additionally, COVID-19 has reinforced the importance of developing vaccine platforms that can be rapidly adapted to respond to future pandemics.

There are different SARS-CoV-2 vaccine candidates at various stages of development and clinical testing, including novel platforms based on DNA or mRNA.^[3]^ At the time of writing, the COVID-19 mRNA vaccines made by Pfizer/BioNTech and Moderna have been approved for emergency use authorization by the FDA. While mRNA vaccines will play a significant role in mitigating effects of the pandemic in areas like the US and parts of Europe, they face manufacturing and distribution limitations that constrain their utility in low-resource settings. Subunit vaccines based on recombinant proteins are more stable and less reliant on the cold chain, making them easier to produce and distribute worldwide.^[4]^ Importantly, large-scale production capacity for subunit vaccines already exists throughout the world.^[5]^ Subunit vaccines are highly customizable and therefore can be very effective in older and immunocompromised populations.^[6]^ COVID-19 will not be contained until people all around the globe are protected from the virus and thus cost, vaccine stability, ease of manufacturing, and ease of distribution are critical qualities to consider during vaccine development.^[4]^

The receptor-binding domain (RBD) of the spike protein that coats the surface of SARS-CoV-2 is an appealing target antigen for COVID-19 subunit vaccines.^[4]^ RBD is the portion of spike that binds to the human angiotensin converting enzyme 2 (ACE2) receptor to mediate viral infection. RBD is more stable than the spike trimer and is manufactured using low-cost, scalable expression platforms.^[7]^ Remarkably, literature reports show that expression levels of RBD can be 100-times greater than expression levels of spike trimer as measured by mass of protein recovered.^[7, 8]^ Considerations to halve COVID-19 vaccine doses (*i*.*e*., only provide one injection when two are recommended) or increase the time between doses highlight the need for greater scalability of vaccines.^[9]^ Additionally, RBD is the target for many neutralizing antibodies that have been identified.^[10]^ An analysis of antibodies produced by survivors of COVID-19 showed that a larger proportion of RBD-binding antibodies were neutralizing compared to antibodies that bound spike outside of the RBD domain.^[11]^ Unfortunately, RBD is poorly immunogenic on its own. In this work we aim to improve the immunogenicity of RBD by prolonging the delivery of the RBD antigen and potent clinically de-risked adjuvants from an injectable hydrogel.

Slow delivery of antigen(s) can result in a more potent humoral immune response. Previous work showed that slow delivery of an HIV vaccine over the course of 2-4 weeks from an implantable osmotic pump led to 20-30 times higher neutralizing antibody titers compared to conventional bolus administration in non-human primates.^[12]^ Microneedle patches are a less-invasive alternative to osmotic pumps, but unfortunately they often require harsh synthesis and loading processes that can damage antigens.^[13]^ Other approaches for slow delivery of antigen have also been pursued, including engineering immunogens that bind to Alum to further improve Alum’s depot effects.^[14]^ As an alternative, our lab and others have developed hydrogel platforms for vaccine delivery that maintain the aqueous solvation of the subunit antigen proteins.^[15]^ Hydrogels are easy to manufacture and can be designed to mimic the mechanics of human tissues.^[16]^ Unfortunately, many hydrogels are covalently cross-linked requiring invasive implantation or complex materials systems that polymerize *in situ*, severely limiting their translatability and compatibility with a range of proteins and other molecules of interest.^[16]^ To address these limitations, many efforts have focused on the development of shear-thinning and self-healing physical hydrogels that can be easily injected and which can afford sustained delivery of molecular cargo.^[17]^ Specifically, our lab has developed an injectable polymer-nanoparticle (PNP) hydrogel system that is extremely inexpensive and scalable and can enable the sustained delivery of a diverse range of vaccine cargo.^[18, 19]^ Roth et al. showed that PNP hydrogel vaccines promote greater affinity maturation and generate durable, robust humoral responses with multiple subunit antigens when leveraging potent adjuvants that have not yet been clinically evaluated.^[18, 20]^ Furthermore, our lab has demonstrated the ability to achieve slow delivery of vaccines through control over PNP hydrogel formulation. Specifically, PNP hydrogels were able to slow down release of both Ovalbumin and Poly(I:C) in a subunit vaccine, resulting in similar release kinetics between both components despite their differences in size and other physicochemical properties.^[21]^ While many adjuvants and combinations of adjuvants have shown great efficacy in pre-clinical models, very few combinations have been evaluated clinically. Additionally, while the versatility of the PNP hydrogel has not been previously demonstrated, the material properties of the PNP hydrogel uniquely enable the comparison of a wide variety of physicochemically distinct adjuvants and their ability to improve the immunogenicity of an antigen with the same platform. We found that a standard bolus injection of the adjuvants Alum (Alhydrogel), AddaVax (an MF59-like squalene emulsion), and CpG + Alum (similar to Dynavax’s CpG/Alum adjuvant) were not sufficient to improve RBD titers after one immunization and were still unable to afford neutralizing responses following both a prime and boost. Of note, Yang et al. observed neutralizing responses following immunization with RBD and Alum in mice, but we were unable to replicate these findings.^[22]^

Here we show that sustained exposure of RBD subunit vaccines comprising clinically de-risked adjuvants within an injectable PNP hydrogel depot increases total anti-RBD IgG titers when compared to the same vaccines administered as a bolus injection. Notably, the high titers elicited by hydrogel-based vaccines are maintained against three common variants of the SARS-CoV-2 Spike protein, Alpha (B.1.1.7), Beta (B.1.351), and Delta (B.1.617.2) whereas the titers elicited by typical bolus vaccines dropped precipitously. Further, a lentiviral SARS-CoV-2 pseudovirus assay revealed that neutralizing responses exceeded those of convalescent human serum after immunization with our hydrogel-based vaccine comprising CpG and Alum. These results suggest that delivery of RBD subunit vaccines in our PNP hydrogel significantly enhances the immunogenicity of RBD and induces neutralizing humoral immunity.

## 2. Results

### Hydrogel for Sustained Vaccine Exposure

RBD was used as the antigen for all vaccine formulations in this work because of its high expression levels, ease of manufacturing, and stability (**Figure 1**a,b). Due to RBD’s small size (~25 kDa; D_h_~5 nm), it exhibits inefficient lymphatic uptake, thereby limiting RBD’s interaction with critical immune cells.^[23]^ Small antigens like RBD often exhibit poor pharmacokinetics because they are quickly dispersed from the injection site and are cleared rapidly.^[24]^ To prolong RBD’s exposure and interaction with immune cells, we sought to further develop an injectable polymer-nanoparticle (PNP) hydrogel that was previously reported by our lab. ^[18, 25]^ An ideal hydrogel depot technology would exhibit several critical mechanical features: (i) mild gelation conditions to enhance vaccine stability in formulation and storage, (ii) high degree of shear thinning for facile injection, (iii) rapid self-healing to mitigate burst release of the vaccine cargo, (iv) sufficiently high yield stress to enable formation of a robust depot that will persist under the normal stresses present in the tissues following administration, and (v) prolonged co-delivery of physicochemically distinct vaccine components. We have demonstrated previously that the PNP hydrogel system satisfies these desired criteria. PNP hydrogels efficiently encapsulate vaccine components without the need for external stimuli, are easily injected using standard syringes and needles,^[26]^ and can co-deliver diverse cargo over prolonged timeframes.^[18]^

**Figure 1.**
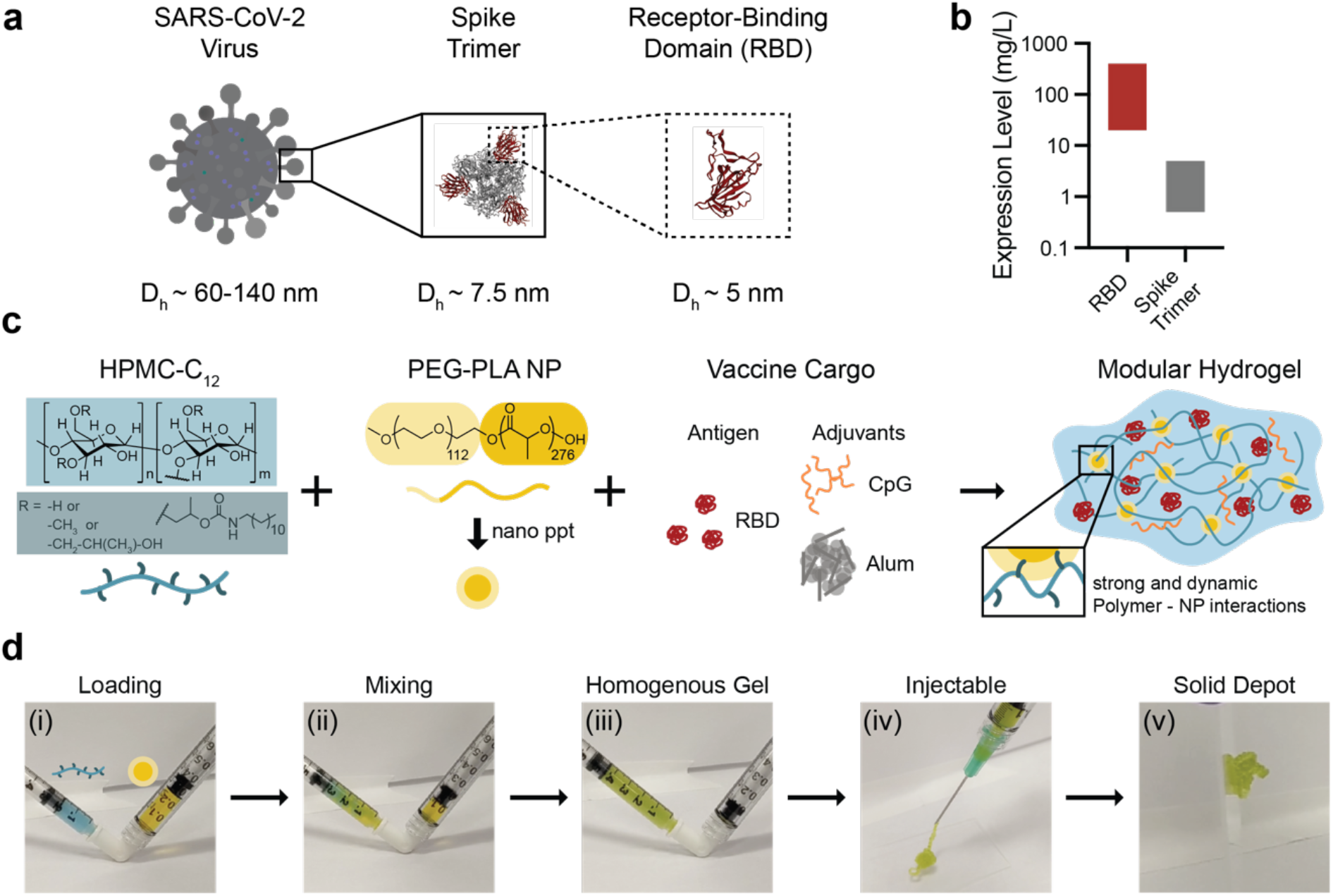
Polymer-nanoparticle (PNP) hydrogel is suitable for subcutaneous delivery of RBD and combinations of clinically de-risked adjuvants. (a) Schematic showing the entire SARS-CoV-2 virus (~60-140 nm), the spike trimer on its surface (~7.5 nm), and the receptor-binding domain (RBD; ~5 nm) that is used as the antigen in these studies. (b) RBD expression levels greatly exceed (~100X) spike trimer expression levels. Bars show the approximate range of expression levels found in the literature.^[8, 24]^ (c) Dodecyl-modified hydroxypropylmethylcellulose (HPMC-C_12_) is combined with poly(ethylene glycol)-*b*- poly(lactic acid) (PEG-PLA) and vaccine cargo (RBD, CpG, and Alum) to form PNP hydrogels. Dynamic, multivalent noncovalent interactions between the polymer and nanoparticles (NPs) leads to physical crosslinking within the hydrogel that behaves like a molecular velcro. (d) HPMC-C_12_ is loaded into one syringe (blue) and the NP solution and vaccine components are loaded into the other (yellow). By connecting the syringes with an elbow (i) and rapidly mixing (ii), a homogenous, solid-like gel is formed (iii). The gel is then easily injected through a 21-guage needle (iv) before self-healing and reforming a solid depot (v) in the subcutaneous space.

PNP hydrogels form rapidly upon mixing hydrophobically-modified hydroxypropylmethylcellulose derivates (HPMC-C_12_) and biodegradable polymeric nanoparticles (NPs) made of poly(ethylene glycol)-*b*-poly(lactic acid) (**Figure 1**c). By adding antigen and adjuvant(s) to the NP solution prior to mixing with the complementary HPMC-C_12_ solution, vaccine components are readily incorporated into the aqueous phase of the hydrogel (**Figure 1**c,d). To prepare PNP hydrogels, an HPMC-C_12_ solution loaded into one syringe and an NP solution comprising the desired vaccine components loaded into another syringe are mixed using an ‘elbow’ connector, forming a homogenous gel (**Figure 1**d). These vaccine-loaded gels are easily injected through a needle before self-healing to form a solid depot at the site of administration (**Figure 1**d; Videos S1,S2).^[26]^

To boost the immunogenicity of these hydrogel-based RBD vaccines, we incorporated combinations of clinically de-risked adjuvants. This work focuses primarily on hydrogel-based vaccines comprising RBD, class B CpG ODN1826 (CpG), and Alhydrogel (Alum) (**Figure 1**d) on account of the broad utility of the CpG/Alum adjuvant system in FDA-approved vaccines (e.g., Heplisav-B) and numerous SARS-CoV-2 subunit vaccine candidates currently in clinical development.^[27]^ Additionally, we leveraged the unique cargo delivery properties of the PNP hydrogel system to directly compare different combinations of toll-like receptor (TLR) and NOD-like receptor (NLR) agonists, including Resiquimod (R848), Monophosphoryl lipid A (MPL), Quil-A saponin (Sap), and the fatty-acid modified form of muramyl dipeptide (MDP) (**Figure S1**).

### Shear-thinning, Self-healing Hydrogel Characterization

We first compared rheological properties of PNP hydrogels both with and without encapsulated Alum to ensure that Alum did not interfere with properties known to be critical for injectability and depot formation.^[18]^ Frequency-dependent oscillatory shear experiments performed in the linear viscoelastic regime showed that the PNP hydrogels with and without Alum had nearly identical frequency responses (**Figure 2**a). For both formulations, the storage modulus (G’) remained above the loss modulus (G”) across the entire range of frequencies evaluated, meaning gels exhibit solid-like properties necessary for robust depot formation (**Figure 2**a).

**Figure 2.**
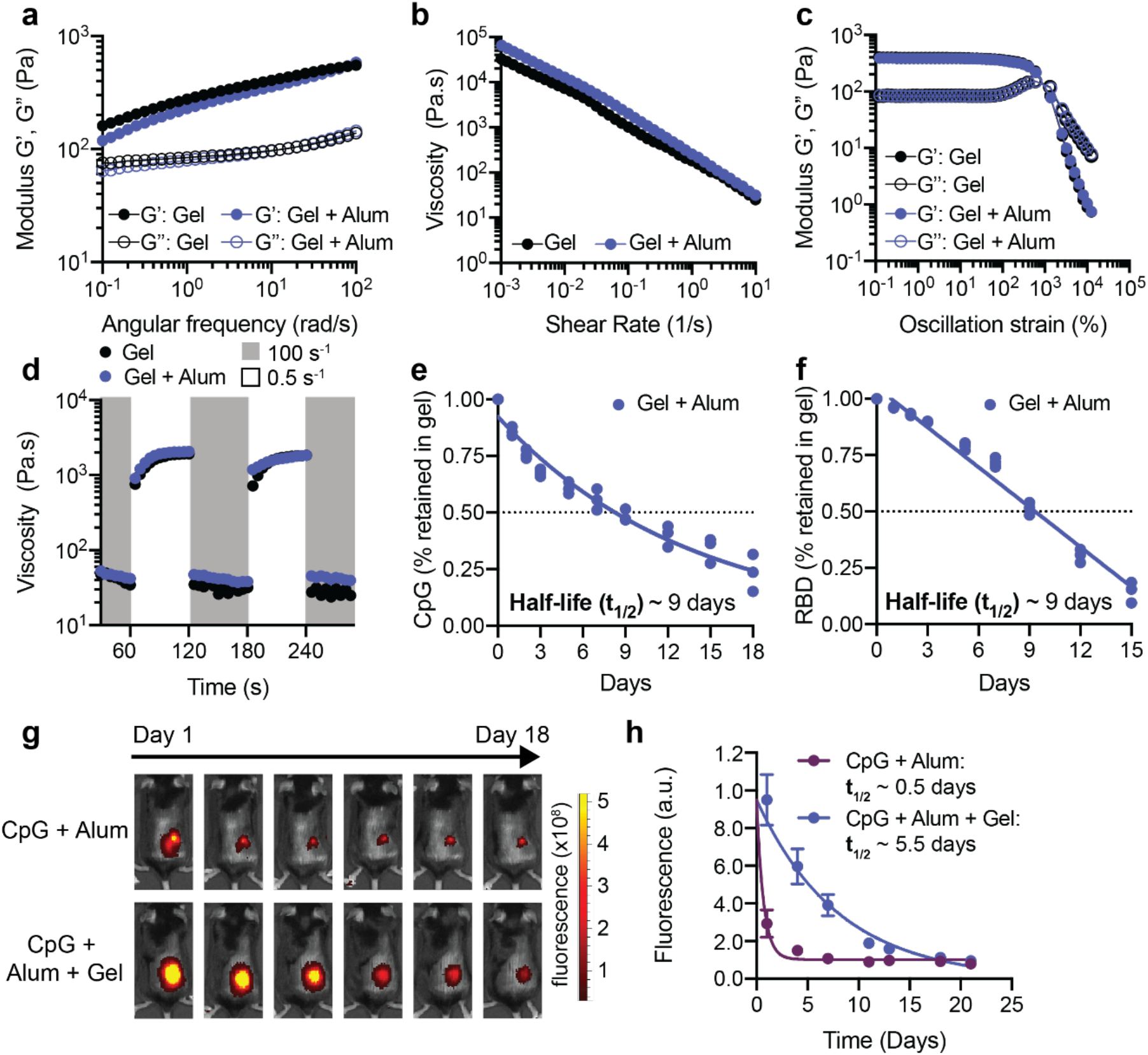
Material properties of PNP hydrogels allow for easy injection, subcutaneous depot formation, and slow release of vaccine cargo. (a) Frequency-dependent oscillatory shear rheology of a PNP hydrogel with or without Alum. (b) Shear-dependent viscosities of PNP hydrogels with or without Alum. (c) Oscillatory amplitude sweeps of PNP hydrogels with or without Alum. The yield stresses were determined by the crossover points and are both around 1300 Pa. (d) Step-shear measurements of hydrogels with or without Alum over three cycles of alternating high shear (gray; 10 s^-1^) and low shear (white; 0.1 s^-1^) rates. (e) Percent of CpG retained in the hydrogel in a glass capillary *in vitro* release study over time. The points were fit with a one-phase decay in GraphPad Prism and the half-life of release was determined. (f) Percent of RBD retained in the same hydrogels as in part e. The points were fit with a linear fit in GraphPad Prism and the half-life of release was determined. (e,f) Each point represents a separate hydrogel (n=3). (g) Representative images demonstrating the different duration of release of Alexa-fluor 647-labeled RBD antigen given as a bolus or gel subcutaneous immunization over 18 days. (h) Fluorescent signal from Alexa-fluor 647-labeled RBD (representative images shown in g) for 3 weeks following immunization as determined by an In Vivo Imaging System (IVIS) (n=5). The points were fit with a one phase-decay in GraphPad Prism and the half-lives were determined. (h) Data are shown as mean +/- SEM.

Hydrogel injectability depends on shear-responsive properties. A shear rate sweep showed that our hydrogel materials are highly shear-thinning, whereby the viscosity of the hydrogels (with or without Alum) decreased several orders of magnitude as the shear rate increased (**Figure 2**b). To assess yielding behavior of the hydrogels, a dynamic amplitude sweep was performed at a frequency of 10 rad/s. For both hydrogels, a yield stress of about 1300 Pa was measured at the crossover point of G’ and G” (**Figure 2**c). Injectability was then tested by measuring the recovery of material properties when alternating between a high shear rate (10 s^-1^) and a low shear rate (0.1 s^-1^) (**Figure 2**d). The viscosity of the hydrogels with and without Alum decreased by about two orders of magnitude under high shear, and rapidly (<5s) recovered when the shear rate was decreased (**Figure 2**d). This test of shear-induced thinning followed by self-healing of the hydrogels mimics an injection through a needle (high shear rate) and the subsequent subcutaneous (SC) depot formation (low shear rate). These data demonstrate that a solid hydrogel depot will form and remain in the SC space after injection which allows for slow release of cargo over time.

### Kinetics of Vaccine Release from the Hydrogel

Previous research has shown that proteins and negatively charged molecules can adsorb to Alum.^[28]^ We hypothesized that the addition of Alum would ensure the RBD protein and the negatively charged CpG would be entrapped within the hydrogel structure to prolong the duration of release. An *in vitro* release assay was used to quantify the kinetics of release of CpG and RBD from the Alum-containing hydrogel over time (**Figure 2**e,f). Additional studies were conducted to quantify kinetics of release of CpG and RBD, as well as other adjuvants (e.g., R848), from the hydrogel without Alum to understand how the physicochemical properties of the adjuvants impacts their release (**Figure S2**a-c). In these assays, the hydrogel was injected into the bottom of a capillary tube and buffer was added above to provide a large sink for release. Tubes were incubated at 37°C to mimic physiological conditions and the entire buffer sink was removed and replaced at each timepoint shown. CpG and RBD were released slowly from the gel containing Alum, with retention half-lives of about nine days (**Figure 2**e,f). The small molecule R848 was also released slowly from the gel with an equivalent half-life of about nine days (**Figure S2**a-c). In gels without Alum, CpG released more rapidly with a half-life of about 2.5 days, while RBD release was unaffected by the absence of Alum (**Figure S3**a-b). These observations indicate that the PNP hydrogel system enables prolonged co-delivery of the RBD antigen and the CpG/Alum adjuvant complex.

Since *in vitro* release studies differ greatly from release within a living organism, we quantified retention of RBD in the hydrogels following SC injection in C57BL/6 mice. RBD was conjugated to an Alexa Fluor 647 dye (AF647-RBD) and administered with CpG and Alum adjuvants to mice via transcutaneous injection in either a gel vehicle or as a bolus injection. The amount of RBD retained in the hydrogel over 18 days was monitored by fluorescence IVIS imaging. Dye-labeled RBD from the Alum-containing bolus treatment was almost undetectable within about a week, while RBD from the hydrogel treatment persisted for the duration of the study, showing that the hydrogel was necessary for prolonged antigen retention (**Figure 2**g). We fit fluorescence values over time with a one-phase exponential decay and determined that the half-life of RBD release was extended from about 0.5 days (95%CI: 0.36 – 0.58) in a standard bolus injection to 5.5 days (95%CI: 3.70 – 9.10) when delivered in the hydrogel (**Figure 2**h). This half-life measured *in vivo* is similar in magnitude to the half-life measured *in vitro* and the difference we observe is likely because cells can actively transport RBD out of the hydrogel *in vivo*.

### Response to Vaccination

To evaluate whether RBD delivery in an adjuvanted hydrogel enhanced the humoral immune response to the antigen, we quantified antigen-specific antibody titers over time in C57BL/6 mice (n=5 each). We were primarily interested in the difference in response to vaccination between the CpG + Alum + Gel group and the dose-matched bolus control group since this comparison allowed us to understand the contribution of sustained vaccine exposure from the hydrogel while keeping the vaccine identity consistent. All immunizations contained 10 µg of RBD. In addition to total IgG antibody titers, we quantified different antibody classes and subclasses (IgM, IgG1, IgG2b, IgG2c) to assess the quality of the response, the anti-spike IgG antibody response, and the acute cytokine response shortly after administration (**Figure 3**a). Vaccines were administered on day 0 and mice were boosted with the original treatment on week 8. Serum was collected weekly, and assays were run at the timepoints shown in **Figure 3a**. High systemic levels of certain cytokines is correlated with toxicity in mice and humans.^[29]^ We measured IFNα and TNFα concentrations at 3-hours post-immunization as an indicator of toxicity for each formulation. The only treatments that led to detectable cytokine levels at 3 hours were R848 + Sap + Gel and R848 + MDP + Gel (**Figure S4**). The IFNα serum concentrations for these treatments were 1-2 ng/mL and the TNFα concentrations were below 0.5 ng/mL. These data suggest that the CpG + Alum + Gel and other treatments that did not include R848 were well-tolerated by this measure.

**Figure 3.**
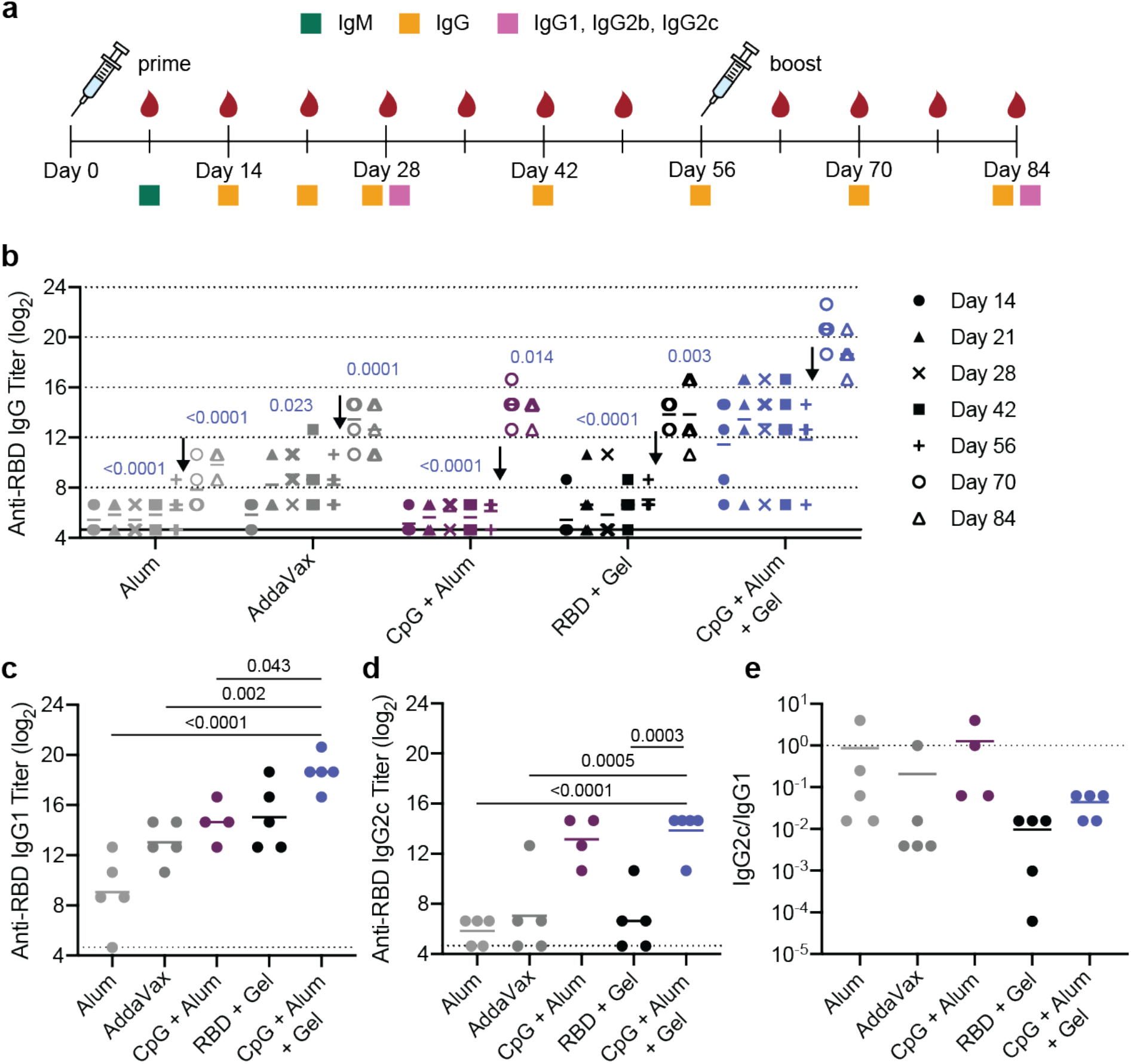
Hydrogel RBD vaccine increases antibody titers compared to bolus vaccine. (a) Timeline of mouse immunizations and blood collection for different assays. Mice were immunized on day 0 and day 56. Serum was collected over time to determine IgG titers. IgM titers were assessed at day 7 (**Figure S**6). IgG1, IgG2b, IgG2c titers were quantified, and neutralization assays were conducted on day 28 and day 84 serum. (b) Anti-RBD IgG ELISA titers before and after boosting (arrow) of several controls and the CpG + Alum + Gel group of interest. P values listed were determined using a 2way ANOVA with Tukey’s multiple comparisons test. P values for comparisons between the CpG + Alum + Gel group and all other groups for day 28 and day 84 are shown above the points. (c-d) Anti-RBD IgG1 (c) and IgG2c (d) titers from serum collected 4 weeks after mice were boosted. P values listed were determined using a one-way ANOVA with Tukey’s multiple comparisons between the CpG + Alum + Gel group and each control group. (e) The ratio of Anti-RBD IgG2c to IgG1 post-boost titers. Lower values (below 1) suggest a Th2 response or skewing towards a stronger humoral response. All data are shown as individual mouse titer values (n=5) and the mean.

Both before and after boosting, mice treated with CpG and Alum in the PNP hydrogel (CpG + Alum + Gel) had higher total antigen-specific IgG antibody titers than Alum, AddaVax, CpG + Alum bolus control, and hydrogel with the RBD antigen only (RBD + Gel) (**Figure 3**b). After boosting, the CpG + Alum + Gel treatment led to titers that were ~60 times greater than all controls, including the bolus treatment that contained identical antigen and adjuvants (**Figure 3**b). As expected, there was a notable increase in titer across all groups following the boost. Additional hydrogel formulations containing RBD and other adjuvant combinations were also evaluated. The hydrogel group loaded with MPL, QuilA, and RBD (MPL + Sap + Gel) resulted in similar titers to the CpG + Alum + Gel treatment (**Figure S5**). This particular adjuvant pair, MPL and QuilA, is utilized in a suspension formulation in the clinical adjuvant system AS01. Notably, although R848-containing gels elicited an increase in serum cytokines at early timepoints, these treatments were not as effective at inducing high antibody titers (**Figure S5**).

IgM is the first antibody isotype produced in response to vaccination prior to class switching.^[30]^ The function of IgM antibodies is to recognize and eliminate pathogens in the early stage of immune defense. On day 7 following immunization, we observed consistent IgM titers across groups (**Figure S6**). Anti-RBD IgG1, IgG2b, and IgG2c titers were determined 4-weeks after both the prime and boost immunizations. RBD-specific IgG1 titers followed a similar trend to total IgG titers (**Figure 3**c; **Figure S7**). Hydrogels comprising CpG and QuilA (CpG + Sap + Gel) led to the highest IgG2b and IgG2c titers (**Figure S7**). CpG + Alum + Gel and CpG + Alum bolus treatments led to higher IgG2b titers than both Alum and AddaVax controls (**Figure S7**d). Although the CpG + Sap + Gel and CpG + Alum + Gel groups maintained high IgG2c titers, the clinically relevant controls (Alum and AddaVax) were much lower (**Figure 3**d, **Figure S7**). The ratio of IgG2c to IgG1 titers is often used as a metric for Th1 versus Th2 skewing.^[31]^ We found that RBD + Gel treatment led to the lowest ratio, suggesting greater Th2 skewing in the humoral response, corroborating previous observations from our lab (**Figure 3**e).^[18]^ The addition of CpG to the hydrogel skewed the ratio slightly more towards a balanced response that was similar in magnitude to what was observed for AddaVax, which is known to promote both strong cellular and humoral immune responses in humans (**Figure 3**e).^[32]^ The Alum and CpG + Alum bolus controls had ratios of about 1, suggesting the most balanced Th2/Th1 response of the groups tested (**Figure 3**e). With the exception of the CpG + Sap hydrogel group, hydrogel treatments tended to skew towards a stronger humoral response, consistent with previous observations (**Figure S8**).^[18]^

Previous work from our lab showed that prolonged germinal center (GC) activity following hydrogel vaccination led to a robust humoral response.^[18]^ We immunized mice with the CpG + Alum + Gel vaccine or the dose-matched bolus control and assessed GC activity at week 2 when we expected the GC response to a bolus immunization should be strongest. We observed no difference in frequency of GC B cells, light zone/dark zone ratio, or frequency of T follicular helper T cells between hydrogel and bolus administrations at this time point, indicating that hydrogel-based delivery does not alter the short-term GC response to the CpG adjuvant (**Figure S9**a-c, **Figure S10**). Next, we sought to compare GC activity over longer time scales to determine if the hydrogel vaccine led to an extended response compared to the bolus vaccine. To do this, we measured the concentration of CXCL13 in serum since it is a biomarker of GC activity that could be quantified from serum from 4-8 weeks post-vaccination (**Figure S9**d-e, **Figure S10**).^[33]^ The median half-life of CXCL13 decay from its peak at week 4 was extended from about 1 week for the bolus vaccine to over 2.5 weeks when the vaccine was delivered from the hydrogel (**Figure S9**f, **Figure S10**).

### SARS-CoV-2 spike-pseudotyped Viral Neutralization Assay

ELISA titers provide a useful measure for understanding antibody binding. In comparison, functional assays like neutralization assays with pseudotyped viruses provide additional information about the humoral response by quantifying antibody-mediated viral inhibition.^[33]^ Recent work by Khoury et al. found that neutralizing antibody levels following vaccination are highly predictive of protection from SARS-CoV-2, similar to what was previously shown in non-human primates.^[35]^ To analyze neutralizing titers, we used lentivirus pseudotyped with SARS-CoV-2 spike and assessed inhibition of viral entry into HeLa cells overexpressing human ACE2.^[11, 34, 36]^ We assessed the neutralizing ability of antibodies in serum collected 4 weeks after the final immunization (week 12). An initial screen with a 1:250 serum dilution showed that the average percent infectivity was reduced more than 50%for the CpG + Alum + Gel, CpG + Sap + Gel, and MPL + Sap + Gel treatments (**Figure S11**). The control groups (Alum and RBD + Gel) showed less significant reductions in infectivity with the exception of AddaVax which showed a slight decrease in average infectivity (~20%). We then ran a series of serum dilutions in experimental duplicate from mice that received CpG + Alum treatments (either gel or bolus) or Alum alone (**Figure 4**a). The CpG + Alum bolus treatment did not notably reduce infectivity even at high serum concentrations (**Figure 4**a). In contrast, serum from mice that received a prime and boost of the CpG + Alum + Gel was completely neutralizing at high concentrations and the IC_50_ could be quantified from dose-inhibition curves from all samples (**Figure 4**a,c).

**Figure 4.**
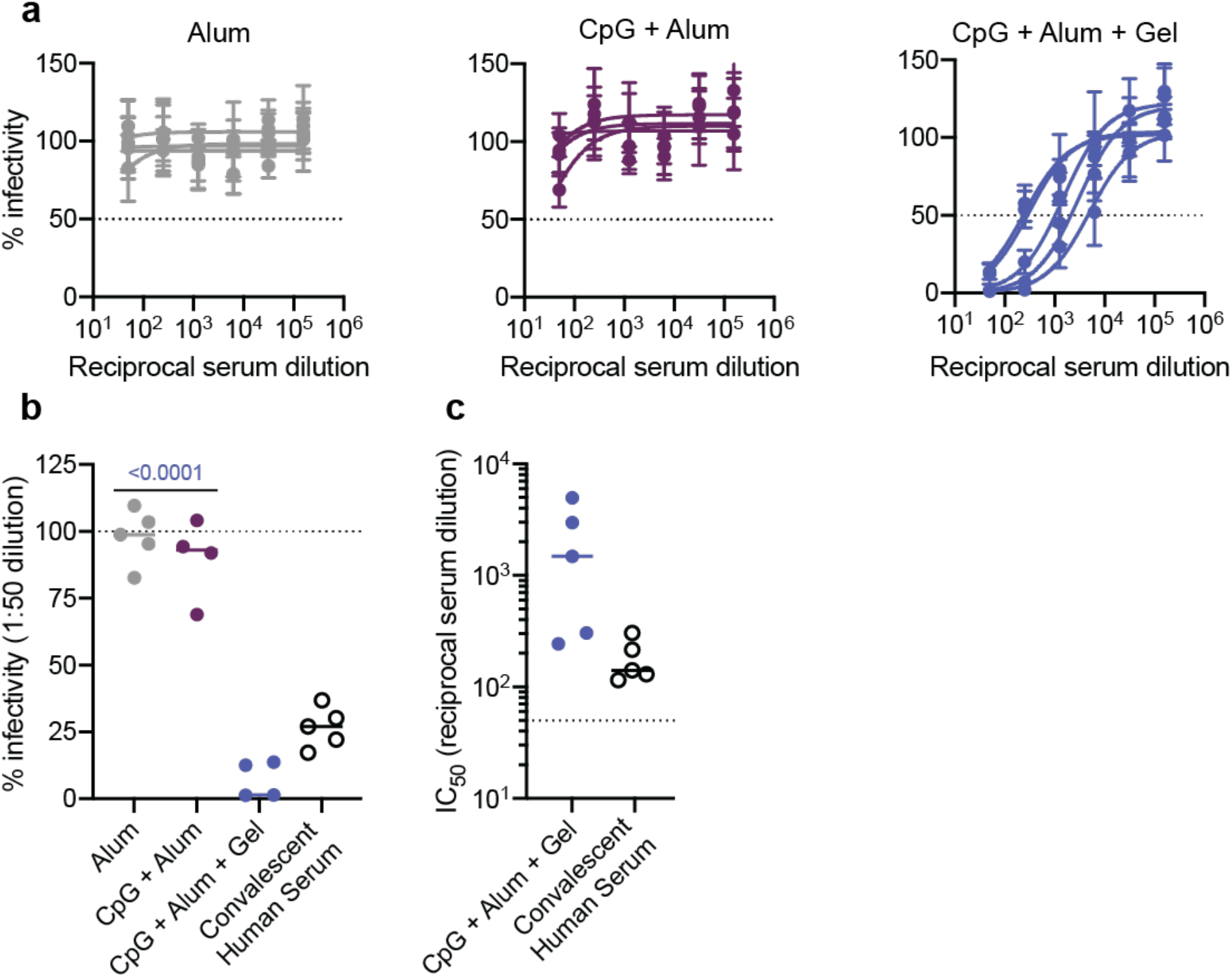
Hydrogel RBD vaccine elicits neutralizing antibodies in mice. (a) Percent infectivity for Alum, CpG + Alum, and CpG + Alum + Gel treatments at a range of serum dilutions as determined by a SARS-CoV-2 spike-pseudotyped viral neutralization assay. Week 12 serum samples were tested for all groups. (b) Percent infectivity for the same treatment groups at a 1 in 50 serum dilution. Convalescent human serum collected 9-10 weeks after the onset of symptoms is also shown for comparison. (c) IC_50_ values determined from the neutralization curves in (a). Samples with neutralizing activity that was undetectable at a 1:50 dilution are excluded. (a) Data shown are mean + /- SEM (n=5). Samples were run in technical duplicate on two separate occasions and values were averaged to determine the mean at each serum dilution. (b-c) Data are shown as individual mouse or human titer values (n=5) and the mean. P values listed were determined in GraphPad Prism software using a one-way ANOVA with Tukey’s multiple comparison test and correspond to comparisons to CpG + Alum + Gel.

By plotting percent infectivity at a 1:50 serum dilution the differences between groups were distinct, with the hydrogel treatment affording greater protection (**Figure 4**b). By this metric, antibodies from convalescent human serum provided similar, but slightly reduced protection compared to antibodies from the hydrogel group (**Figure 4**b). Overall, quantifiable levels of neutralizing antibodies were observed in all mice that received two immunizations of CpG + Alum + Gel (**Figure 4**c). No neutralization was detected in samples from mice that received either Alum or the CpG + Alum bolus treatment so IC_50_ values could not be determined (**Figure 4**c). Markedly, the CpG + Alum + Gel prime/boost treatment led to antibodies with a mean IC_50_ value that was about an order of magnitude greater than the mean IC_50_ value from antibodies in the convalescent human serum samples (**Figure 4**c).

Several different SARS-CoV-2 variants of concern with increased transmission rates have recently been identified in countries around the world and given Greek-letter identifiers.^[37]^ Studies have been conducted to determine if previous infection and/or immunization with current vaccines protects against these variants. Although the Moderna and Pfizer/BioNTech are thought to provide robust protection against the Alpha (B.1.1.7) variant, results vary against the Beta (B.1.351) variant.^[37]^ Neutralizing antibodies from COVID-19 patients have been identified that bind regions of RBD that are conserved across emerging variants of SARS-CoV-2 and across other coronaviruses, signifying that RBD might be a useful target antigen for achieving broad antibody responses.^[38]^ Unfortunately, SARS-CoV-2 will continue to mutate until protective immunizations are distributed equitably around the globe. Here we assessed titers against the native spike as well as the Alpha (B.1.1.7), Beta (B.1.351), and Delta (B.1.617.2) variant spike proteins following immunization with our vaccines to see if broad protection resulted.

Overall, native wildtype spike titers reflected similar trends to those observed with anti-RBD IgG titers (**Figure 5**a, **Figure S5**), but were slightly lower than anti-RBD titers as expected since we vaccinated with RBD. All vaccines led to similar titers against the Alpha (B.1.1.7) variant and the wildtype form, but titers against the Beta (B.1.351) and Delta (B.1.617.2) variants were noticeably lower compared to the wildtype across all groups except Alum and CpG + Alum + Gel (**Figure 5**a). In particular, the fold reduction in mean Beta (B.1.351) variant titers compared to mean wildtype titers was about 7.0 for the bolus CpG + Alum group and only about 1.7 for the comparable hydrogel, indicating that the hydrogel vaccine provided broader coverage against this highly evasive variant of concern (**Figure 5**a). Similarly, a fold reduction in mean Delta (B.1.617.2) variant titers of 2.8 was observed for the bolus CpG + Alum group and only about 1.2 for the comparable hydrogel. The broader coverage provided by the hydrogel vaccine against these variants is particularly notable since multiple key mutations in the Beta (B.1.351) variant, including the N501Y mutation linked to increased viral transmission, as well as mutations in the Delta (B1.617.2) variant, exist within the RBD (**Figure 5**b).

**Figure 5.**
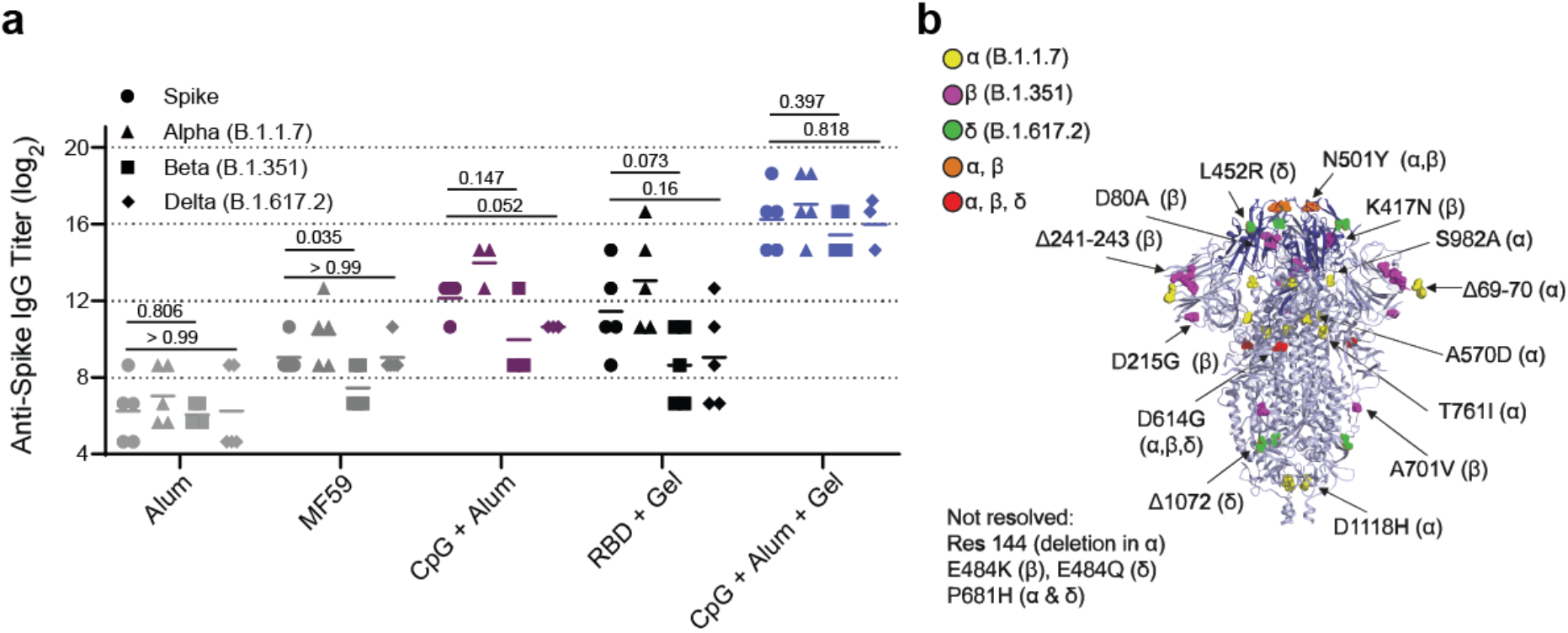
Hydrogel RBD vaccine provides a more potent and broader response against spike and spike variants. (a) Anti-spike IgG ELISA titers from serum collected 4 weeks after the final immunization (Day 84). Titers were determined for wildtype spike as well as Alpha (B.1.1.7), Beta (B.1.351), and Delta (B.1.617.2) variants of spike. P values correspond to t tests comparing anti-spike titers between wildtype vs. Beta (B.1.351) variant of spike and wildtype vs. Delta (B.1.617.2) variant. (b) SARS-CoV-2 spike trimer with highlighted mutations for the Alpha (B.1.1.7), Beta (B.1.351), and Delta (B.1.617.2) variants. RBD is shown in darker blue and corresponding bolded mutations exist within the RBD region.^[38]^

### Single-Injection Hydrogel Vaccine Induces a Durable Response

An ideal COVID-19 vaccine would require a single immunization. We therefore conducted a preliminary study using a single-injection vaccine based on the CpG + Alum + Gel treatment that contained double the dose of all components (2X Gel) with the goal of inducing similarly high titers and neutralization (**Figure S12**a). IgG titers from the 2X Gel vaccine exceeded those of post-prime single dose gel titers and exceeded the post-boost titers of the bolus control (**Figure S12**b). Without boosting, the 2X Gel IgG titers persisted for 12 weeks. The anti-RBD IgG titers also remained above the titers of convalescent human serum showing that the vaccine efficacy in mice surpasses immunity following a natural infection in humans. Strikingly, the anti-spike IgG 2X Gel titers were almost equivalent to the post-boost CpG + Alum + Gel titers, suggesting a single shot achieved the same humoral response to the native spike protein (**Figure S12**c). The anti-spike titers for the 2X Gel also persisted and remained above the anti-spike titers for the bolus group through week 12 and remained at or above the titer levels of the convalescent human patient serum (**Figure S12**c).

To further compare the response following 2X Gel treatment to the prime-boost responses, we also measured 2X Gel IgG1 and IgG2c titers. 2X Gel titers generally followed a similar trend to the CpG + Alum + Gel titers with higher mean IgG1 titers and lower mean IgG2c titers compared to the bolus (**Figure S12**d,e). The IgG2c/IgG1 ratio was similar to that of the CpG + Alum + Gel group, suggesting a Th2-skewed response as expected (**Figure S12**f). We also evaluated the 2X gel in the spike-pseudotyped neutralization assay and found that serum from all mice exhibited neutralizing responses, but that the effect was slightly weaker than what was observed following the prime/boost hydrogel immunizations (**Figure S13**a-c).

## 3. Discussion

In this work we improved the immunogenicity of RBD by delivering it in an injectable, slow-release hydrogel formulated with two clinically de-risked adjuvants, CpG and Alum. The PNP hydrogel platform is compatible with many physiochemically distinct molecules, which allowed us to screen a variety of adjuvant combinations before further pursuing CpG and Alum. Additionally, since a primary benefit of RBD as an antigen is its superior stability, we have evaluated RBD stability upon encapsulation within the PNP hydrogel and confirmed that the RBD antigen completely maintains its integrity in the hydrogel formulation for weeks under constant mechanical agitation at 50°C (**Figure S14**). The prime/boost CpG + Alum + Gel vaccine led to a significant increase in titers against both RBD and spike compared to the bolus control and vaccines containing other common adjuvants, Alum and AddaVax. Notably, the prime/boost vaccine induced neutralizing antibody titers of approximately 10^3^ reciprocal serum dilution, which is significantly greater than the IC_50_ of the bolus control and the convalescent human serum tested. It is important to highlight that accelerated Th1 responses in humans are correlated with less severe COVID-19 disease, whereas strongly Th2-skewed responses following vaccination with inactivated SARS-CoV viral vaccines are associated with enhancement in lung disease.^[40]^ The CpG + Alum + Gel immunization led to overall well balanced Th1 and Th2 responses, and the potent neutralizing responses observed for this vaccine highlights that there is no antibody-dependent enhancement of infectivity. Our hydrogel vaccine also resulted in similarly high titers against Alpha (B.1.1.7), Beta (B.1.351), and Delta (B.1.617.2) spike variants, suggesting broad protection against newly emerging, highly infectious strains.

To address the need for single-injection vaccines, we conducted a preliminary study with a single hydrogel vaccine administration comprising the same total antigen and adjuvant dose, which we called “2X Gel”. Following a single immunization with the 2X Gel, titers were greater than the those of the prime/boost bolus group and nearly reached the level of the prime/boost hydrogel vaccine group. Similar titers to prime/boost bolus treatment also resulted when testing titers against the Alpha (B.1.1.7), Beta (B.1.351), and Delta (B.1.617.2) variants. Future experiments should be conducted to determine if our single injection hydrogel vaccine can provide more robust neutralization and long-term protection.

It is also important to highlight that we focus on humoral immune responses in this study as antibody levels, and particularly neutralizing antibody levels, have become recognized as strong correlates of protection against severe COVID-19.^[35]^ Our current work does not yet address cellular immunity such as T cell-mediated responses, which have also been shown to offer long-term and broad protection against SARS-CoV-2 variants in convalescent individuals.^[41]^ Future work will evaluate T cell-mediated immunity induced by our PNP hydrogel vaccines.

At the time of writing, the Pfizer/BioNTech and Moderna mRNA vaccines have been approved for emergency use by the FDA.^[42]^ While this is exciting news, it is unlikely that these vaccines, or other mRNA vaccines, will reach large parts of the globe. Currently, there are only a few manufacturing sites for the mRNA vaccines, and they are concentrated in the US, Belgium, and Germany. Both vaccines require frozen storage and the Pfizer/BioNTech vaccine must be kept around -70°C, drastically limiting distribution due to the significant logistical challenges of maintaining this cold-chain.^[42, 43]^ It is therefore necessary to continue developing vaccines that are more stable, less reliant on the cold-chain, and that can be manufactured at more sites world-wide. To this end, we pursued a subunit vaccine with RBD as the antigen since it is easy to manufacture, very stable, and therefore recommended for the development of low-cost, accessible COVID-19 vaccines.^[4]^ Additionally, we focused our efforts on adjuvants that are already approved for other uses or are advanced in the clinical pipeline. Another class B CpG, CpG 1018, was developed by Dynavax and is currently being used in collaborations with the Coalition for Epidemic Preparedness Innovations (CEPI), the University of Queensland, and Clover Biopharmaceuticals to augment various COVID-19 vaccines.^[44]^

In addition to the choice of antigen and adjuvant, there is a growing need for novel and efficient drug delivery systems like our PNP hydrogel platform.^[45]^ The hydrogel is injectable and forms a depot that allows for slow release of the vaccine over several days to weeks, which is an order of magnitude longer than the duration of release from Alum alone (**Figure 2**g,h). Ultimately, this leads to greater affinity maturation and a more potent and durable humoral response.^[18]^ In this work we validated that, unlike other hydrogel or microneedle vaccine platforms, the PNP hydrogel is readily loaded with diverse molecular cargo, allowing for the antigen and adjuvants to be co-presented to immune cells. In addition, we found that delivery of the CpG + Alum RBD subunit vaccine in our hydrogel improved breadth of coverage against emerging SARS-CoV-2 spike variants as compared to the vaccine-matched bolus control. As discussed in previous work from our lab, the hydrogel likely acts through two main mechanisms: (i) as a local stimulatory niche where infiltrating cells experience high local concentrations of adjuvant and antigen and (ii) as a slow-release depot that allows for prolonged vaccine exposure. As a local stimulatory niche, vaccine-loaded hydrogels increase recruitment of antigen presenting cells such as macrophages and dendritic cells that crucial to initiating the adaptive immune response.^[18]^ While we observe an increase in immunogenicity with our current hydrogel-based RBD subunit vaccine comprising a CpG/Alum adjuvant complex, future investigations will aim to characterize immune cell infiltration into the hydrogel and the impact of the formation of such a local inflammatory niche on downstream humoral immune responses. By delivering RBD with the clinically de-risked adjuvants, CpG and Alum, in our PNP hydrogel, we were able to achieve titers and neutralization levels that greatly exceeded those of the relevant clinical controls, Alum and AddaVax, as well as the bolus CpG + Alum vaccine.

## 4. Methods

### Materials

HPMC (meets USP testing specifications), N,N-diisopropylethylamine (Hunig’s base), hexanes, diethyl ether, N-methyl-2-pyrrolidone (NMP), dichloromethane (DCM), lactide (LA), 1-dodecylisocynate, and diazobicylcoundecene (DBU) were purchased from Sigma-Aldrich and used as received. Monomethoxy-PEG (5 kDa) was purchased from Sigma-Aldrich and was purified by azeotropic distillation with toluene prior to use.

### Preparation of HPMC-C_12_

HPMC−C_12_ was prepared according to previously reported procedures.^22, 30, 52^ HPMC (1.0 g) was dissolved in NMP (40 mL) by stirring at 80°C for 1 h. Once the solution reached room temperature (RT), 1-dodecylisocynate (105 mg, 0.5 mmol) and N,N-diisopropylethylamine (catalyst, ∼3 drops) were dissolved in NMP (5.0 mL). This solution was added dropwise to the reaction mixture, which was then stirred at RT for 16 h. This solution was then precipitated from acetone, decanted, redissolved in water (∼2 wt %), and placed in a dialysis tube for dialysis for 3−4 days. The polymer was lyophilized and reconstituted to a 60 mg/mL solution with sterile PBS.

### Preparation of PEG−PLA NPs

PEG−PLA was prepared as previously reported.^18, 25^ Monomethoxy-PEG (5 kDa; 0.25 g, 4.1 mmol) and DBU (15 µL, 0.1 mmol; 1.4 mol %relative to LA) were dissolved in anhydrous dichloromethane (1.0 mL). LA (1.0 g, 6.9 mmol) was dissolved in anhydrous DCM (3.0 mL) with mild heating. The LA solution was added rapidly to the PEG/DBU solution and was allowed to stir for 10 min. The reaction mixture was quenched and precipitated by a 1:1 hexane and ethyl ether solution. The synthesized PEG−PLA was collected and dried under vacuum. Gel permeation chromatography (GPC) was used to verify that the molecular weight and dispersity of polymers meet our quality control (QC) parameters. NPs were prepared as previously reported.^18, 25^ A 1 mL solution of PEG−PLA in DMSO (50 mg/mL) was added dropwise to 10 mL of water at RT under a high stir rate (600 rpm). NPs were purified by centrifugation over a filter (molecular weight cutoff of 10 kDa; Millipore Amicon Ultra-15) followed by resuspension in PBS to a final concentration of 200 mg/mL. NPs were characterized by dynamic light scattering (DLS) to find the NP diameter, 37 ± 4 nm.

### PNP Hydrogel Preparation

The hydrogel formulation contained 2 wt %HPMC−C_12_ and 10 wt %PEG−PLA NPs in PBS. These gels were made by mixing a 2:3:1 weight ratio of 6 wt %HPMC−C12 polymer solution, 20 wt %NP solution, and PBS containing all other vaccine components. The NP and aqueous components were loaded into one syringe, the HPMC-C_12_ was loaded into a second syringe and components were mixed using an elbow connector. After mixing, the elbow was replaced with a 21-gauge needle for injection.

### Material Characterization

Rheological characterization was performed on PNP hydrogels with or without Alum (Alhydrogel) using a TA Instruments Discovery HR-2 torque-controlled rheometer (TA Instruments) fitted with a Peltier stage. All measurements were performed using a serrated 20 mm plate geometry at 25°C with a 700 µm gap height. Dynamic oscillatory frequency sweep measurements were performed from 0.1 to 100 rad/s with a constant oscillation strain in the linear viscoelastic regime (1%). Amplitude sweeps were performed at a constant angular frequency of 10 rad/s from 0.01%to 10,000%strain with a gap height of 500 µm. Steady shear experiments were performed by alternating between a low shear rate (0.1 s^-1^) and high shear rate (10 s^-1^) for 60 seconds each for three full cycles. Shear rate sweep experiments were performed from 10 s^-1^ to 0.001 s^-1^.

### Expression and Purification of RBD

The mammalian expression plasmid for RBD production was a kind give from Dr. Florian Krammer and was previously described in detail in (Amanat et al, 2020, Nat Medicine). RBD was expressed and purified from Expi293F cells as previously described.^43^ Briefly, Expi293F cells were cultured using 66%FreeStyle293 Expression /33%Expi293 Expression medium (Thermo Fisher) and grown in polycarbonate baffled shaking flasks at 37°C and 8%CO2 while shaking. Cells were transfected at a density of approximately 3-4 × 10^6^ cells/mL. Cells were harvested 3-5 days post-transfection via centrifugation. RBD was purified with HisPur NiNTA resin (Thermo Fisher). Resin/supernatant mixtures were added to glass chromatography columns for gravity flow purification. Resin was washed with 10 mM imidazole/1X PBS [pH 7.4] and proteins were eluted. NiNTA elutions were concentrated using Amicon spin concentrators (10 kDa MWCO for RBD) followed by size-exclusion chromatography. The RBD was purified using a GE Superdex 200 Increase 10/300 GL column. Fractions were pooled based on A280 signals and/or SDS-PAGE. Samples for immunizations were supplemented with 10%glycerol, filtered through a 0.22-µm filter, snap frozen, and stored at -20°C until use.

### Vaccine Formulations

The vaccines contained a 10 µg dose of RBD and combinations of 5 µg Quil-A Adjuvant (Invivogen), 50 µg Resiquimod (R848; Selleck Chemicals), 20 µg L18-MDP (Invivogen), 10 µg MPLA (Invivogen), 20 µg CpG ODN 1826 (Invivogen), or 100 µg Alhydrogel in 100 µL hydrogel or PBS based on the treatment group. For the bolus vaccines, the above vaccine doses were prepared in PBS and loaded into a syringe for administration. For the PNP hydrogels, the vaccine cargo was added at the appropriate concentration into the PBS component of the gel and combined with the NP solution before mixing with the HPMC-C_12_ polymer, as described above.

### RBD and CpG Gel Release Studies

Hydrogels were prepared the same way as described in the “PNP Hydrogel Preparation” section and were loaded with 10 µg RBD, 20 µg CpG and 100 µg Alum. Glass capillary tubes were plugged at one end with epoxy and 100 µL of gel was injected into the bottom of 3 different tubes. 350 µL of PBS was then added on top of each gel. The tubes were stored upright in an incubator at 37°C for about 3 weeks. At each timepoint, ~300 µL of PBS was removed and the same amount was replaced. The amount of RBD released at each timepoint was determined using a Micro BCA™ Protein Assay Kit (Fisher Scientific) following the manufacturer’s instructions (including using the Bovine Serum Albumin standards provided in the kit). The amount of CpG released was determined by measuring the absorbance at 260, subtracting the absorbance from a blank well with buffer, and then applying the Beer-Lambert law with an extinction coefficient of 0.027 µg/mL* cm^-1^ for single-stranded DNA. For both types of cargo, the cumulative release was calculated and normalized to the total amount released over the duration of the experiment. For CpG retention, the points were fit with a one phase-decay in GraphPad Prism and the half-life of release was determined (n=3). For RBD retention, the points were fit with a linear fit in GraphPad Prism and the half-life of release was determined (n=3).

Alexa Fluor 647-conjugated RBD was synthesized by the following methods: AFDye 647-NHS ester (Click Chemistry Tools, 1.8 mg, 1.85 µmol) was added to a solution of RBD protein (0.84 mg, 0.926 µmol) in PBS. The NHS ester reaction was conducted with a 20 molar excess of AFDye 647-NHS ester to RBD in the dark for 3 h at RT with mild shaking. The solution was quenched by diluting 10-fold with PBS. The solution was then purified in centrifugal filters (Amicon Ultra, MWCO 10 kDa) at 4500 RCF for 20 min, and the purification step was repeated until all excess dye was removed.

Vaccine-loaded hydrogels or bolus controls with 10 µg Alexa Fluor 647-conjugated RBD, 20 µg CpG and 100 µg Alum were injected into mice and fluorescence was monitored over time by In Vivo Imaging System (IVIS Lumina Imager; Ex = 600 nm, Em = 670 nm). Images were collected on days 1, 4, 7, 11, 13, and 18 (n=5 mice). Signal was quantified as raw fluorescence within a constant region of interest. GraphPad Prism was used to fit one-phase decays with a constrained initial value based on day 0 signal and half-lives of release were determined (n=5).

### RBD Stressed Aging Studies

Hydrogels were prepared the same way as described in the “PNP Hydrogel Preparation” section and were loaded with 10 µg RBD. After the mixing step, the vaccine-loaded hydrogels or bolus controls were placed in a 50°C incubator on top of a shaker at 200 rpm to agitate. At each timepoint, 20 µl of the bolus samples were extracted and mixed with PBS at a 1:50 dilution for analysis with ELISA. Similarly, 20 µl of the gel samples were extracted and mixed with PBS in two syringes at a 1:50 dilution. For analysis with ELISA, 96-well Maxisorp plates (Thermo Fisher) were coated with 50 µl of the diluted samples overnight at 4°C. Plates were then blocked with 1%bovine serum albumin (BSA in 1X PBS) for 1 h at RT. Spike RBD antibody (Sino Biological 40592-MP01) was added at a 1:2000 dilution and incubated on blocked plates for 2 h at RT. IgG Fc-HRP goat-anti-mouse secondary antibody (Invitrogen A16084) was then added at a 1:10,000 dilution (in 1%BSA) for 1h at RT. Plates were developed with TMB substrate (TMB ELISA Substrate (High Sensitivity), Abcam). The reaction was stopped with 1 M HCl. Plates were analyzed using a Synergy H1 Microplate Reader (BioTek Instruments) at 450 nm. Data was normalized to day 0 timepoint for gel and bolus groups, respectively.

### Mice and Vaccination

C57BL/6 mice were purchased from Charles River and housed at Stanford University. 8-10 week-old female mice were used. Mice were shaved prior to initial immunization. Mice received 100 µL hydrogel or bolus vaccine on their backs under brief isoflurane anesthesia. Bolus treatments were injected with a 26-gauge needle and hydrogels were injected with a 21-gauge needle. Mouse blood was collected from the tail vein for survival bleeds over the course of the study.

### Mouse Serum ELISAs

Anti-RBD and Anti-spike trimer antibody titers were measured using an end-point ELISA. 96-well Maxisorp plates (Thermo Fisher) were coated with RBD, full-length spike^43^, the mutant spike from the Alpha B.1.1.7 (Sino Biological 40591-V08H12), the mutant spike from Beta B.1.351 (Sino Biological 40591-V08H10), or the mutant spike from Delta B.1.617.2 (Sino Biological 40591-V08H23) at 2 µg/mL in 1X PBS [pH 7.4] overnight at 4°C. Plates were then blocked with 1%bovine serum albumin (BSA in 1X PBS) for 1 h at RT. Serum samples were serially diluted starting at a 1:100 dilution and incubated on blocked plates for 2 h at RT. One of the following goat-anti-mouse secondary antibodies was used: IgG Fc-HRP (1:10,000, Invitrogen A16084), IgG1 heavy chain HRP (1:50,000, abcam ab97240), IgG2b heavy chain HRP (1:10,000, abcam ab97250), IgG2c heavy chain HRP (1:10,000, abcam ab97255), or IgM mu chain HRP (1:10,000 abcam ab97230). The secondary antibody was added at the dilution listed (in 1%BSA) for 1 h at RT. 5X PBS-T washes were done between each incubation step. Plates were developed with TMB substrate (TMB ELISA Substrate (High Sensitivity), Abcam). The reaction was stopped with 1 M HCl. Plates were analyzed using a Synergy H1 Microplate Reader (BioTek Instruments) at 450 nm. End-point titers were defined as the highest serum dilution that gave an optical density above 0.1. For plots displaying a single time point, P values listed were determined using a one-way ANOVA or Student’s t-test and for plots displaying multiple timepoints, P values listed were determined using a 2way ANOVA. Both statistical analyses were done using Tukey’s multiple comparisons test on GraphPad Prism software. All titer data is shown as the mean and individual points (n=5) with P values listed above the points.

Mouse IFNa All Subtype ELISA kit, High Sensitivity (PBL Assay Science, 42115-1), Mouse TNFa Quantikine ELISA kit (R&D Systems, SMTA00B), and Legend Max Mouse CXCL13 (BLC) ELISA kit (BioLegend, 441907) were used to quantify different serum cytokines. Serum dilutions of 1:10 were used for all ELISAs. Concentrations were determined by ELISA according to manufacturer’s instructions. Absorbance was measured at 450 nm in a Synergy H1 Microplate Reader (BioTek). Cytokine concentrations were calculated from the standard curves which were run in technical duplicate. Concentration data are reported as ng/mL for IFNa and TNFa and pg/mL for CXCL13 and displayed as individual points and the mean.

### Immunophenotyping in Lymph Nodes

Methods from previous germinal center phenotyping done in the lab were followed.^23^ Briefly, inguinal lymph nodes were removed from mice after euthanasia and were disrupted to create a cell suspension. For flow cytometry analysis, cells were blocked with anti-CD16/CD38 (clone: 2.4G2) and then stained with fluorochrome conjugated antibodies: CD19, GL7, CD95, CXCR4, CD86, IgG1, CD4, CXCR5, and PD1. Cells were then washed, fixed, and analyzed on an LSRII flow cytometer. Data were analyzed with FlowJo 10 (FlowJo LLC). See Table S1 for the antibody panel.

### SARS-CoV-2 spike-pseudotyped Viral Neutralization Assay

Neutralization assays were conducted as described previously.^45^ Briefly, SARS-CoV-2 spike-pseudotyped lentivirus was produced in HEK239T cells. Six million cells were seeded one day prior to transfection. A five-plasmid system was used for viral production.^42^ Plasmids were added to filter-sterilized water and HEPES-buffered saline was added dropwise to a final volume of 1 mL. CaCl_2_ was added dropwise while the solution was agitated to form transfection complexes. Transfection reactions were incubated for 20 min at RT, then added to plated cells. Virus-containing culture supernatants were harvested ~72 hours after transfection by centrifugation and filtered through a 0.45-µm syringe filter. Stocks were stored at -80°C.

For the neutralization assay, ACE2/HeLa cells were plated 1-2 days prior to infection.^14^ Mouse serum was heat inactivated at 56 °C for 30 min prior to use. Mouse serum and virus were diluted in cell culture medium and supplemented with a polybrene at a final concentration of 5 µg/mL. Serum/virus dilutions were incubated at 37 °C for 1 h. After incubation, media was removed from cells and replaced with serum/virus dilutions and incubated at 37°C for 2 days. Cells were then lysed using BriteLite (Perkin Elmer) luciferase readout reagent, and luminescence was measured with a BioTek plate reader. Each plate was normalized by wells with cells only or virus only and curves were fit with a three-parameter non-linear regression inhibitor curve to obtain IC_50_ values. Serum samples that failed to neutralize or that neutralized at levels higher than 1:50 were set at the limit of quantitation for analyses. Serum dilution curves display mean infectivity +/- SEM for each individual mouse (n=5) at each serum dilution. Normalized values were fit with a three-parameter non-linear regression inhibitor curve in GraphPad Prism to obtain IC_50_ values. Fits were constrained to have a value of 0%at the bottom of the fit. Single dilution infectivity plots and IC_50_ data are shown as individual mouse or human titer values (n=5) and the mean. P values listed were determined in GraphPad Prism software using a one-way ANOVA with Tukey’s multiple comparison test.

### Animal Protocol

All animal studies were performed in accordance with National Institutes of Health guidelines and with the approval of the Stanford Administrative Panel on Laboratory Animal Care (APLAC-32109).

### Collection of Serum from Human Patients

Convalescent COVID-19 blood was collected from 5 donors 9-10 weeks after onset of symptoms. Blood was collected in microtubes with serum gel for clotting (Starstedt), centrifuged for 5 minutes at 10,000g and then serum was stored at - 80°C until used. Blood collection was done by finger-prick and was performed in accordance with National Institutes of Health guidelines with the approval of the Stanford Human Subjects Research and IRB Compliance Office (IRB-58511).

## Data Availability Statement

The data that support the findings of this study are available on request from the corresponding author. The data are not publicly available due to privacy or ethical restrictions.

## Acknowledgements

We would like to thank all members of the Appel lab for their useful discussion and advice throughout this project, specifically Catie Meis for sharing her capillary release design. Also, the staff of the BioE/ChemE Animal Facility who cared for our mice, and the Cochran lab for providing access to their In Vivo Imaging System. This research was financially supported by the Center for Human Systems Immunology with the Bill and Melinda Gates Foundation (OPP1113682; OPP1211043). This work was also supported by the Stanford Maternal and Child Health Research Institute postdoctoral fellowship (to AEP) and the Chan Zuckerberg Biohub (to PSK). We thank Dr. Jesse Bloom, Kate Crawford, Dr. Dennis Burton, and Dr. Deli Huang for sharing the plasmids, cells, and invaluable advice for implementation of the spike-pseudotyped lentiviral neutralization assay (to AEP). We thank the NIH Cell and Molecular Biology Training Program (T32 GM007276; to ECG), the Schmidt Science Fellows program, in partnership with the Rhodes Trust (to AID), the National Science Foundation Graduate Research Fellowship (to JY and AKG), the Gabilan Fellowship of the Stanford Graduate Fellowship in Science and Engineering (to AKG), the National Science Foundation under award ECCS-1542152 (to GAR), the Eastman Kodak Fellowship (BSO), and the Stanford NIH T32 Biotechnology Training Grant (5T32GM008412-25; to ELM).

## Author Contributions

ECG, GAR and EAA designed the broad concepts and research studies. AEP, PSK, JA, and BP designed specific experiments. AEP conducted experiments and provided reagents. ECG, GAR, AEP, BSO, ELM, VCTMP, AKG, AID, JY and JA performed research and experiments. ECG and EAA wrote the paper. AEP, ELM, JY and GAR edited the paper.

