## Supplementary Information for "Hydrogel-based slow release of a receptor-binding domain subunit vaccine elicits neutralizing antibody responses against SARS-CoV-2"

**Table of Contents**

Supplemental Figures
Figure S1| Gel synthesis schematic for adjuvant screen……………………………….…..2
Figure S2| R848 and CpG *in vitro* release……………………………………………….3
Figure S3| Serum cytokine levels for adjuvant screen……………………………….…..4
Figure S4| Anti-RBD and Anti-spike titers for adjuvant screen…………………………5
Figure S5| Anti-RBD IgM titers for adjuvant screen…………………………………....6
Figure S6| Anti-RBD titer subtypes for adjuvant screen……………………………….…7
Figure S7| Ratio of IgG2c to IgG1 titers for adjuvant screen……………………………8
Figure S8| Germinal center activity analysis……………………………………….….…9
Figure S9| Sample gating strategy………………………..…………………….…….…10
Figure S10| Adjuvant screen infectivity from neutralization assay…………...………..11
Figure S11| Single injection hydrogel vaccine titers……………………………..……12
Figure S12| Single injection hydrogel neutralization assay……………………………13
Figure S13| 2X hydrogel neutralization assay…………………………………………14
Figure S14| RBD stressed aging assay…………………………………………………15
Supplemental Table
Table S1| Flow cytometry antibody information…………………….……………….....16
Supplemental Videos
Video S1| PNP hydrogel mixing…………………………………………………………17
Video S2| PNP hydrogel injection………………………………………………………..17

### Supplemental Figures

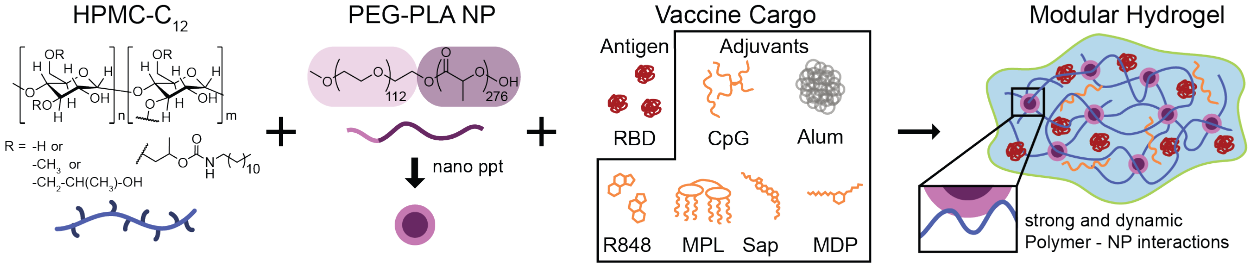


**Figure S1.** HPMC-C12 is combined with PEG-PLA and vaccine cargo to form PNP hydrogels. Dynamic, multivalent noncovalent interactions between the polymer and NPs leads to physical crosslinking within the hydrogel that behaves like molecular velcro. For these studies, different combinations of class B CpG ODN1826 (CpG), Alhydrogel (Alum), Resiquimod (R848), Monophosphoryl lipid A (MPL), Quil-A (Sap), and the fatty-acid modified form of muramyl dipeptide (MDP) were used as adjuvants alongside the RBD antigen.

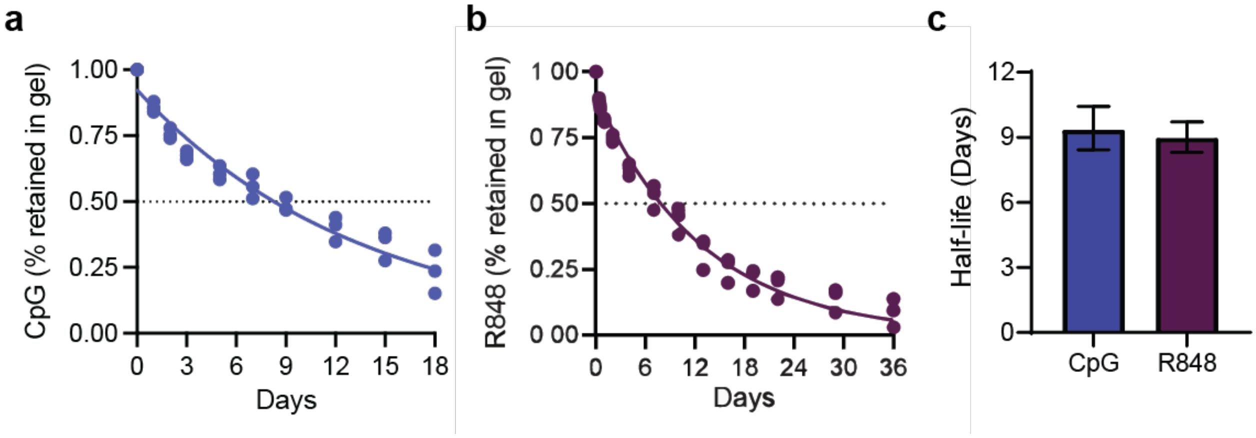


**Figure S2.** CpG (a) and R848 (b) retention in hydrogels over time as determined by a glass capillary *in vitro* release study. The hydrogel containing CpG also contained Alum. Points were fit with a one-phase decay in GraphPad Prism and the half-life of release was determined. Each point represents a separate hydrogel (n=3). (c) Half-lives determined from (a) and (b). The error bar shows the 95% confidence interval.

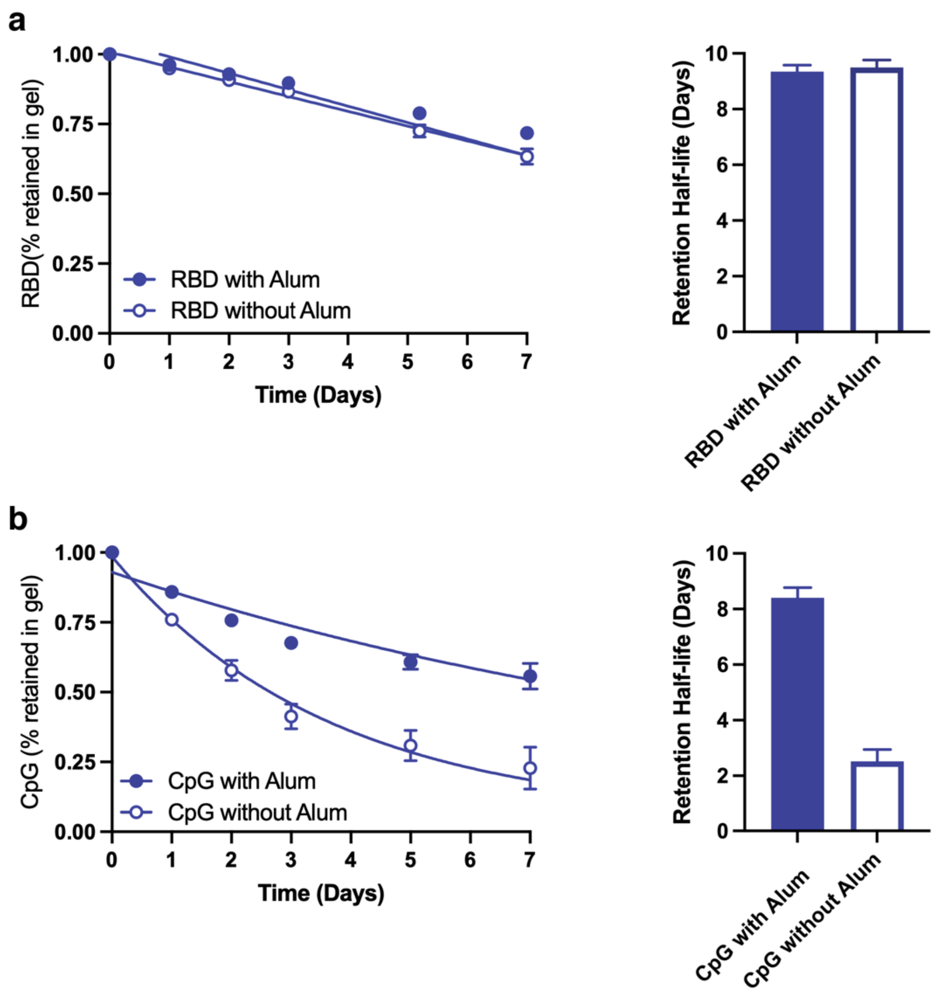


**Figure S3.** RBD (a) and CpG (b) retention in hydrogels with or without Alum over time as determined by a glass capillary *in vitro* release study. Each point represents a separate hydrogel. Points from release curves (left) were fit with either a linear or one-phase decay fit in GraphPad Prism and the half-life of release was determined and reported (right). Data shown as mean ± s.d. (n=3).

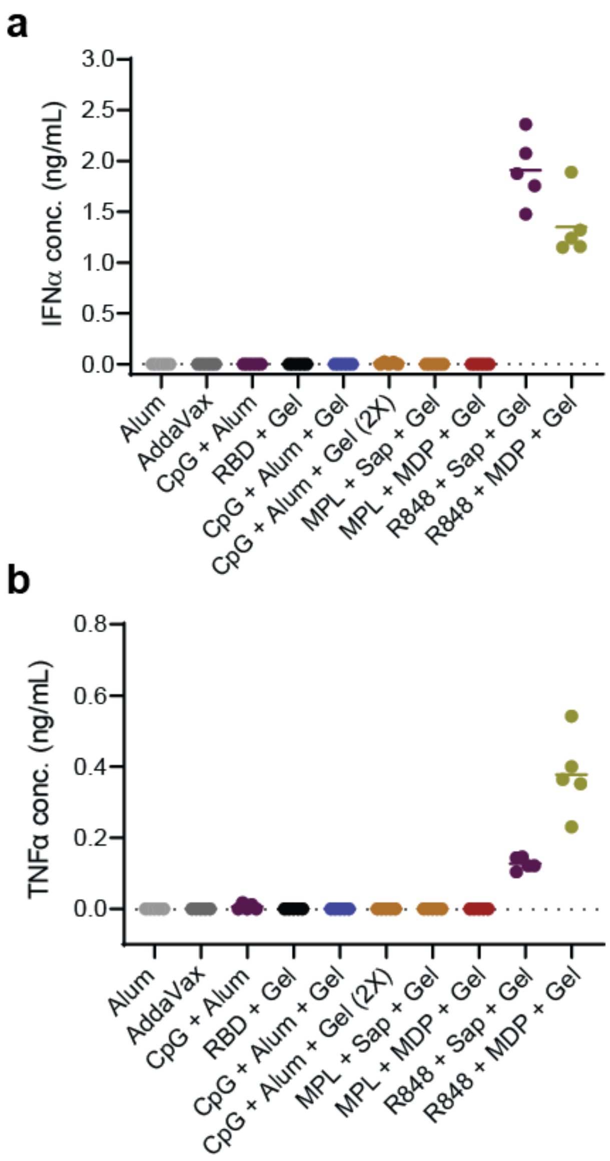


**Figure S4.** IFNα (a) and TNFα (b) levels in serum collected 3 hours after the initial immunization as determined by ELISA (n=5). All treatments contain the adjuvants listed as well as the RBD antigen. The dotted lines show the detection limits of the assays. Individual values that each represent data from a single mouse are shown along with the mean for each group.

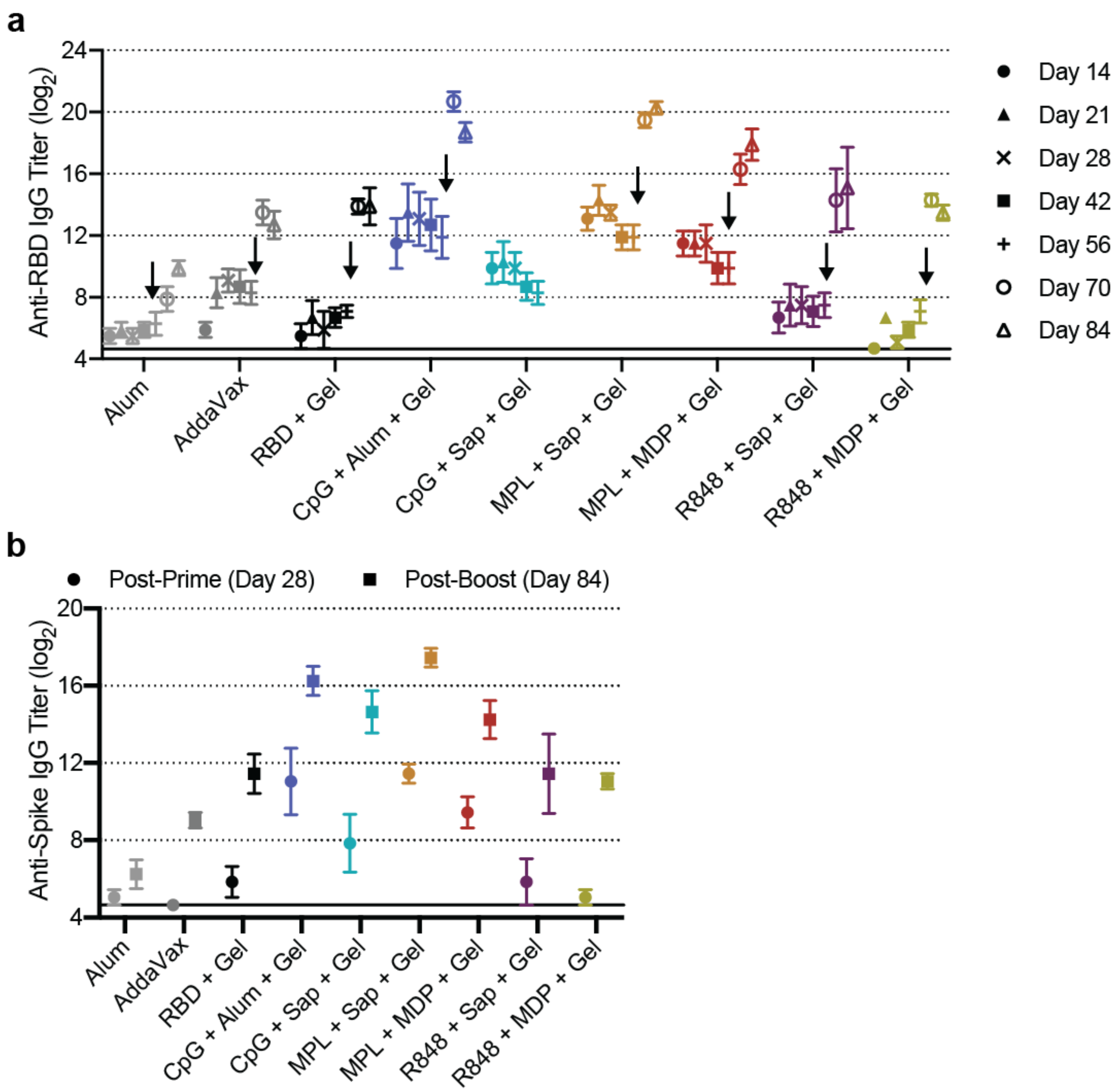


**Figure S5.** Anti-RBD (a) and anti-Spike (b) IgG titers for all hydrogel vaccines tested. All vaccines include the adjuvants listed and the RBD antigen. The arrow shows the point at which mice were given a boost immunization. All data are shown as the mean +/- SEM (n=5).

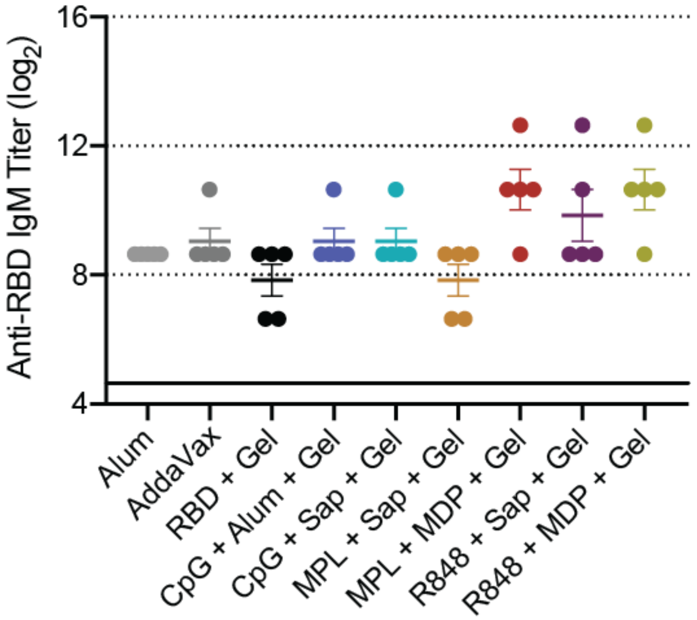


**Figure S6.** Anti-RBD IgM titers of serum collected 7 days after immunization with each of the hydrogel vaccines tested. All vaccines include the adjuvants listed and the RBD antigen. All data are shown as the mean +/- SEM (n=5).

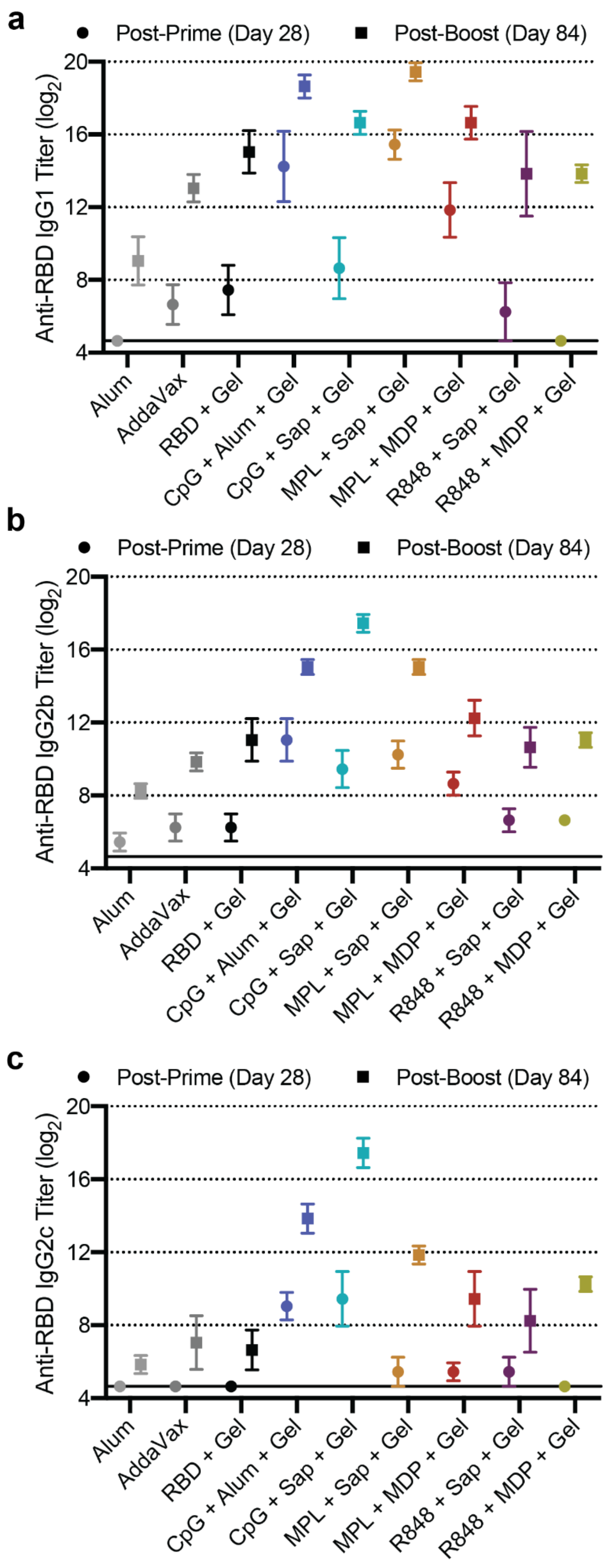


**Figure S7.** Anti-RBD IgG1 (a), IgG2b (b), and IgG2c (c) titers 4 weeks after the prime (Day 28) and the boost (Day 84). All data are shown as the mean +/- SEM (n=5).

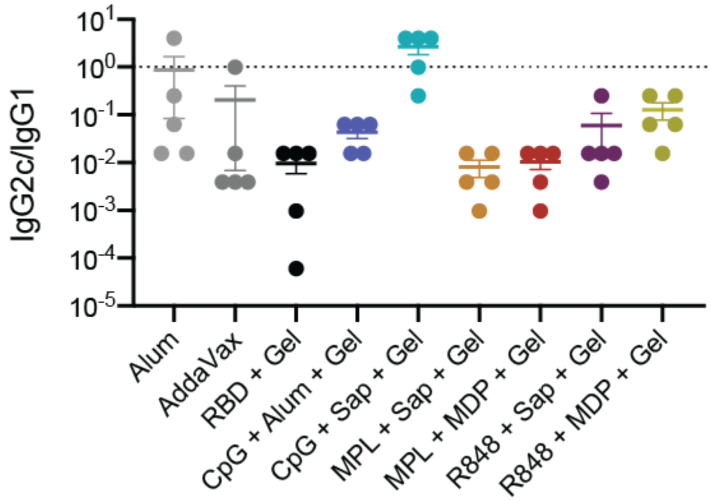


**Figure S8.** The ratio of anti-RBD IgG2c to IgG1 post-boost (Day 84) titers. A value less than one indicates Th2 skewing and a stronger humoral response whereas a value over one indicates a stronger Th1 or cell-mediate response. All data are shown as the mean +/- SEM (n=5).

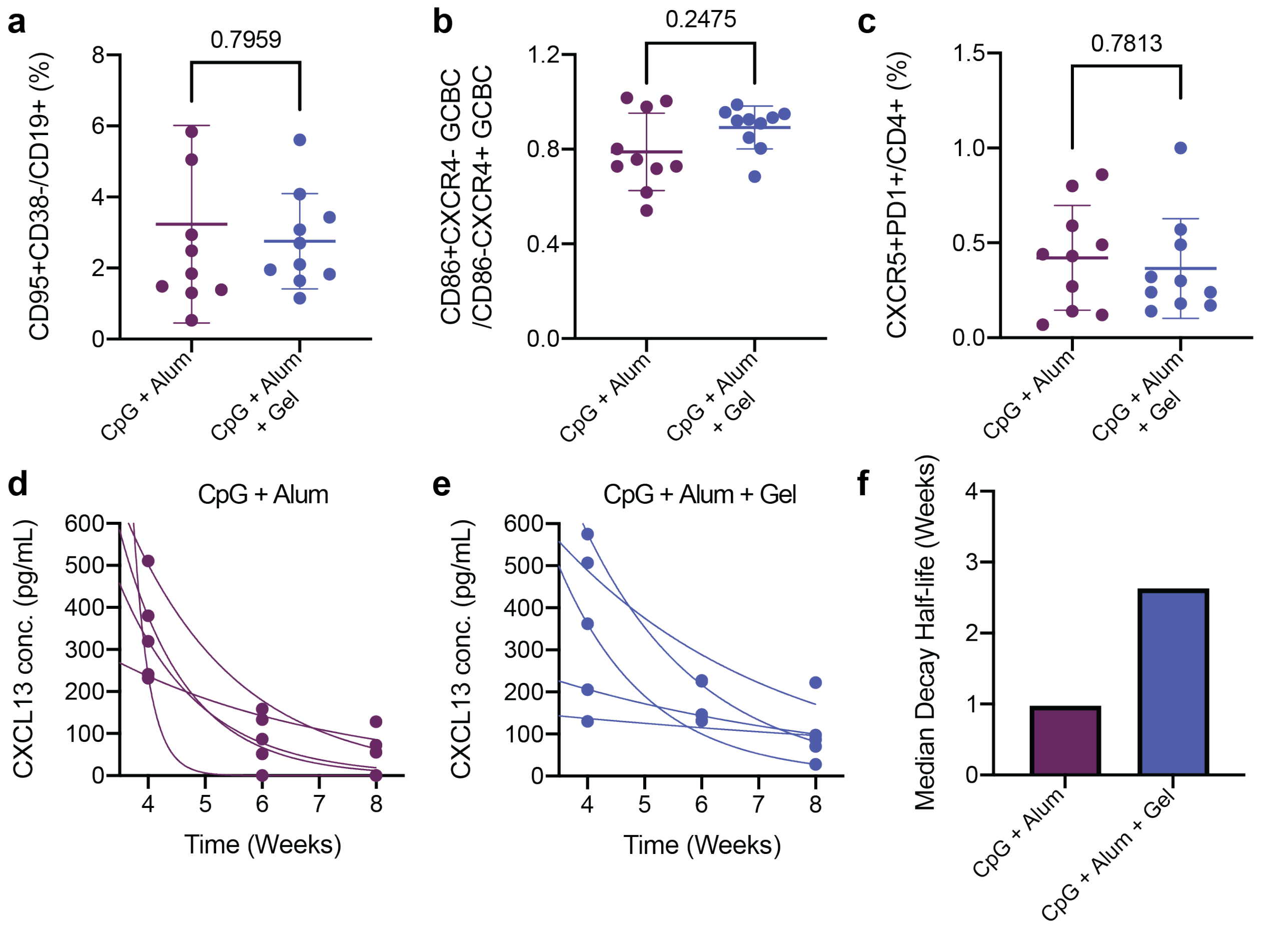


**Figure S9.** Inguinal lymph nodes were collected from mice 2 weeks after immunization. Samples were processed and stained prior to analysis by flow cytometry. (a) Frequency of GC B cells from CD19+ cells following hydrogel or bolus vaccination. (b) Ratio of light zone (LZ) to dark zone (DZ) GC B cells. (c) Frequency of T follicular helper cells from CD4+ T cells. (a-c) All data are shown as mean +/- SEM (n=10) and P values from Mann-Whitney tests are shown. (d-e) CXCL13 concentration from serum collected 4, 6, and 8 weeks after immunization with CpG + Alum (d) or CpG + Alum + Gel (e). Points were fit with a one-phase exponential decay on GraphPad Prism with a lower constraint set to 0. Each curve represents one mouse. (f) Median half-life of decay from CXCL13 peak at week 4. Sample gating strategy is shown in Figure S10.

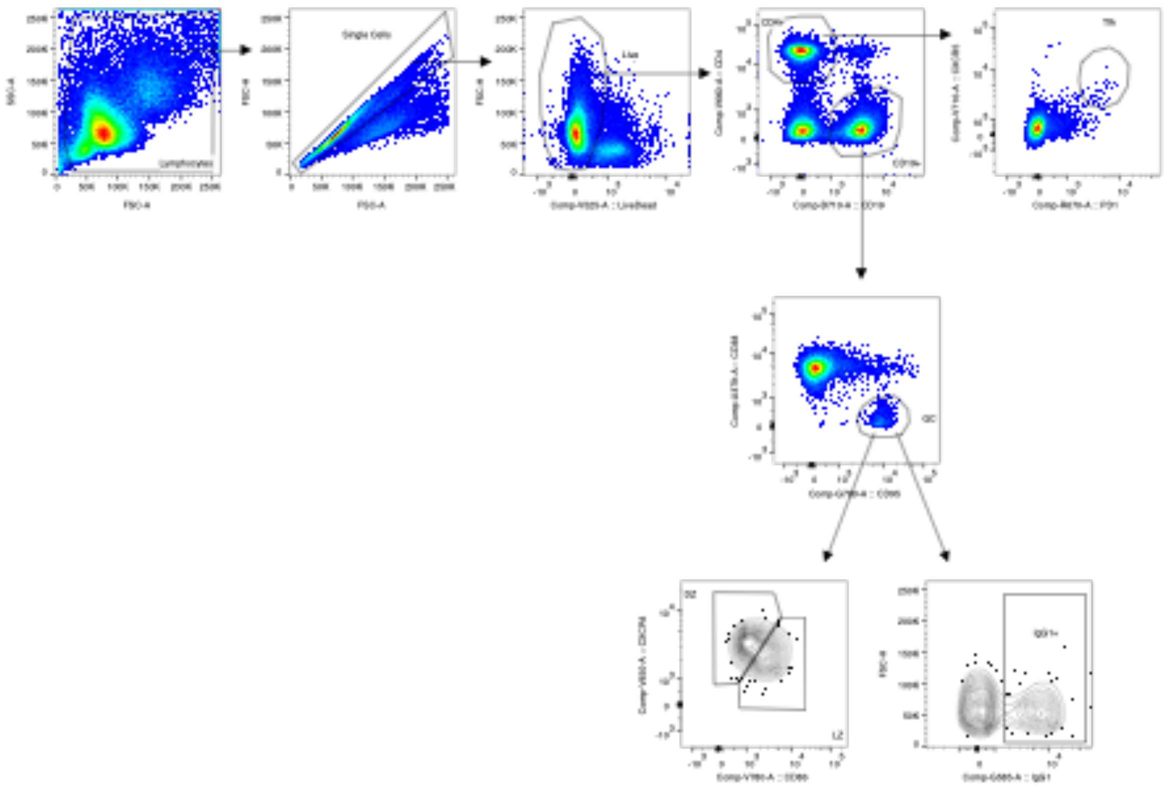


**Figure S10.** Sample gating strategy for plots in Figure S7. Table S1 lists all fluorophores used for gating and analysis.

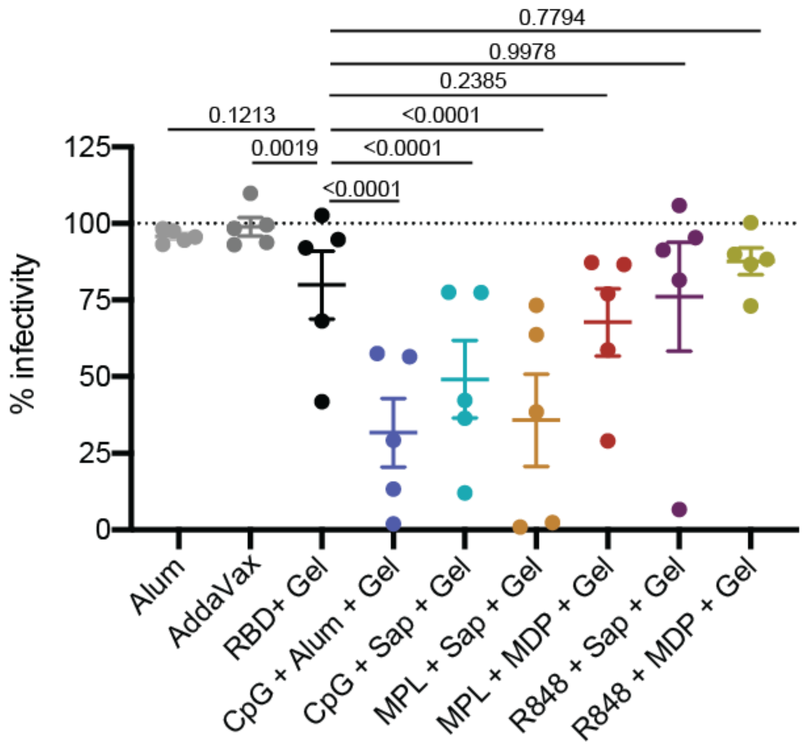


**Figure S11.** Percent infectivity as determined by a SARS-CoV-2 spike-pseudotyped viral neutralization assay at a 1:250 serum dilution. Serum was collected 4 weeks after the boost immunization (Day 84). All points are the average of four normalized infectivity values obtained from two experimental replicates and data are shown as the cohort mean +/- SEM (n=5). P values listed were determined using a 2way ANOVA with Tukey's multiple comparisons test on GraphPad Prism software.

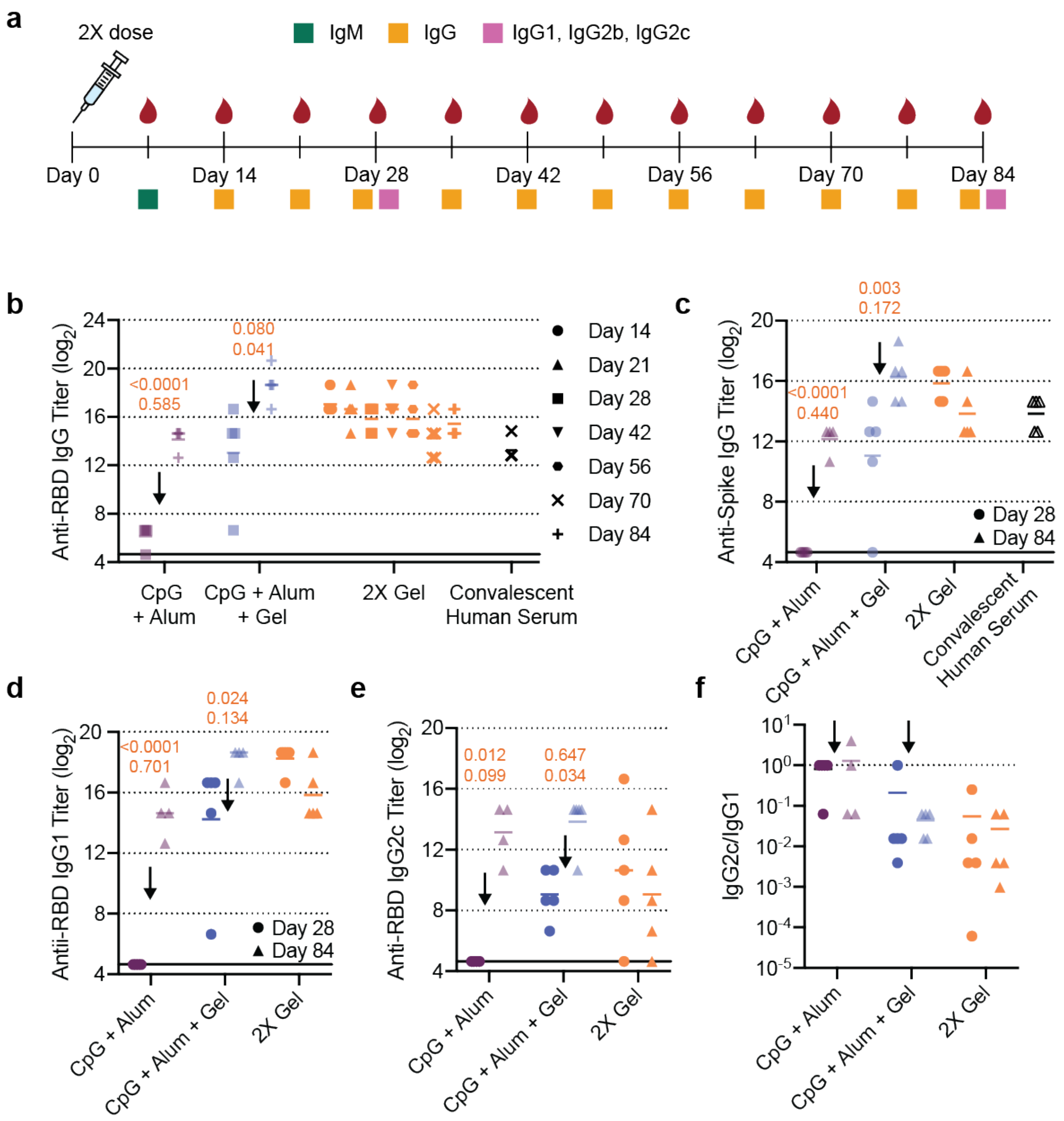


**Figure S12.** (a) Timeline of mouse immunization and blood collection for different assays. The double-dose hydrogel (2X Gel) was administered a single time and no boost was given. (b) Anti-RBD IgG ELISA titers over time. CpG + Alum and CpG + Alum + Gel groups were boosted at week 8, but the 2X Gel group was not. Convalescent human serum is shown for comparison. (c) Anti-spike IgG ELISA titers from day 28 and 84 serum. (d-e) Anti-RBD IgG1 (d) and IgG2c (e) titers from serum collected at day 28 and day 84. (f) Ratio of IgG2c to IgG1 titers where lower values (below 1) suggest a Th2 response or skewing towards a stronger humoral response. Arrows separate pre- and post-boost data. (b-e) P values listed were determined using a 2way ANOVA with Tukey's multiple comparisons test on GraphPad Prism software. P values for comparisons between the 2X Gel group and all other groups for day 28 (top) and day 84 (bottom) are shown above the points. Data points are transparent if they are also shown on a previous figure. All data are shown as individual mouse or human titer values (n=5) and the mean.

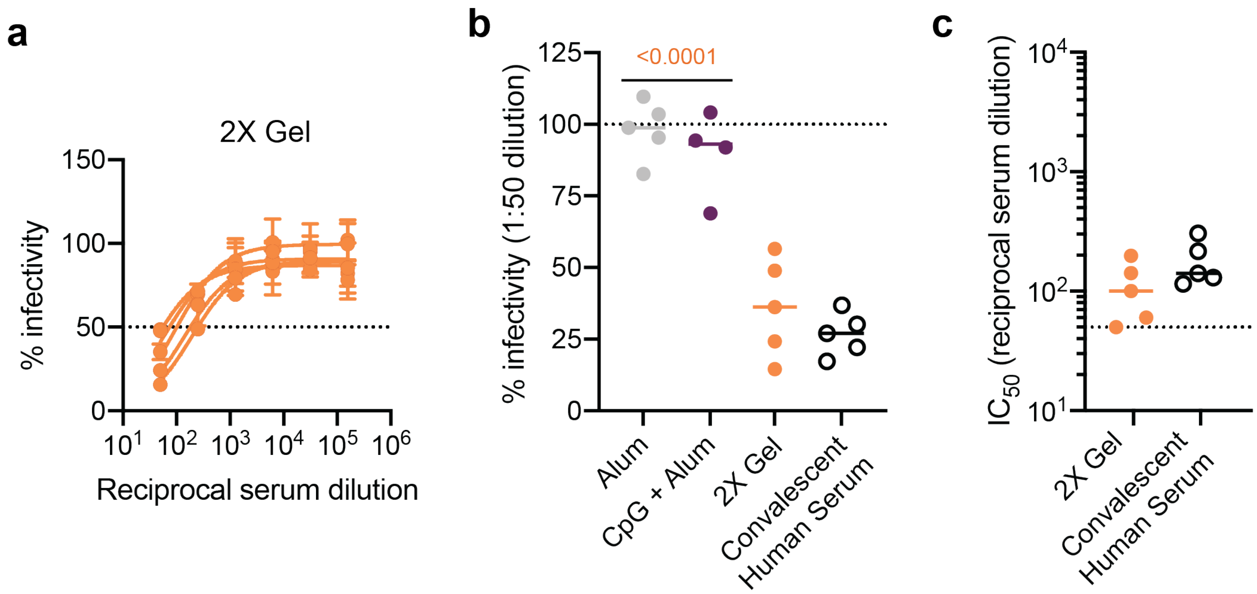


**Figure S13.** (a) Percent infectivity for 2X Gel treatment at a range of serum dilutions as determined by a SARS-CoV-2 spike-pseudotyped viral neutralization assay. Week 4 serum samples were tested. (b) Percent infectivity at a 1 in 50 serum dilution. Neutralizing titers of convalescent human serum collected 9-10 weeks after the onset of symptoms is also shown for comparison. (c) $IC_{50}$ values determined from the neutralization curves in (a). (a) Data are shown mean +/- SEM (n=5). Samples were run in technical duplicate on two separate occasions and values were averaged to determine the mean at each serum dilution. (b-c) Data are shown as individual mouse or human titer values (n=5) and the mean. P values listed were determined in GraphPad Prism software using a one-way ANOVA with Tukey's multiple comparison test. (b) P values corresponding to comparisons to 2X Gel are shown.

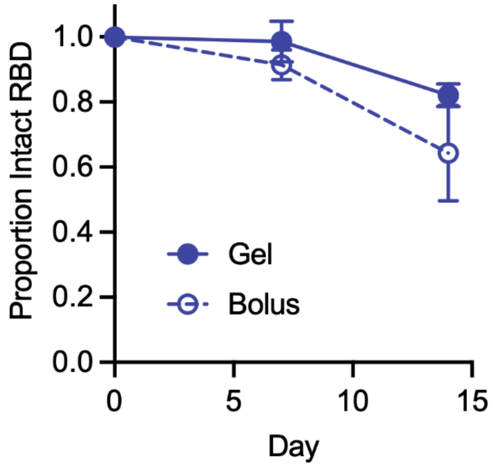


**Figure S14.** Proportion of intact RBD as determined by ELISA at each timepoint over a two-week period after constant mechanical agitation in 50°C. All points are the average of normalized values obtained from experimental replicates and data are shown as the cohort mean +/- s.d. (n=2).

**Table S1.** Flow cytometry antibody information

| Antibody (all anti-mouse) | Manufacturer | Clone |
|---|---|---|
| CD19-PerCP-Cy5.5 | BioLegend | 6D5 |
| GL7-A488 | BioLegend | GL7 |
| CD95-PE-Cy7 | BD Biosciences | Jo2 |
| CXCR4-BV421 | BioLegend | L276F12 |
| CD86-BV785 | Prepared in Pulendran Lab | GL1 |
| IgG1-PE | BD Biosciences | A85-1 |
| CD4-BV650 | BioLegend | GK1.5 |
| CXCR5-BV711 | BioLegend | L138D7 |
| PD1-A647 | BioLegend | 29F.1A12 |

**Supplemental Videos**

**Video S1.** Video of PNP hydrogel mixing. HPMC-$C_{12}$ is loaded in one syringe (blue) and the nanoparticles and vaccine components are loaded into another (yellow). The syringes are attached with an elbow and plungers are pressed back and forth to mix. A homogenous green gel results with the vaccine components, polymer, and nanoparticles evenly distributed throughout the gel.

**Video S2.** Video of PNP hydrogel being injected through a 21-gauge needle. The hydrogel exhibits high viscosity before injection, but dramatically shear-thins and flows through the needle during injection, and rapidly returns to a high viscosity, shape-persistent state once exiting the needle.
